# No Assembly Required: Using BTyper3 to Assess the Congruency of a Proposed Taxonomic Framework for the *Bacillus cereus* group with Historical Typing Methods

**DOI:** 10.1101/2020.06.28.175992

**Authors:** Laura M. Carroll, Rachel A. Cheng, Jasna Kovac

**Affiliations:** Structural and Computational Biology Unit, EMBL, Heidelberg, Germany; Department of Food Science, Cornell University, Ithaca, NY, USA; Department of Food Science, The Pennsylvania State University, University Park, PA, USA

**Keywords:** *Bacillus anthracis*, *Bacillus cereus*, *Bacillus cereus* group, *Bacillus thuringiensis*, foodborne illness, taxonomy

## Abstract

The *Bacillus cereus* group, also known as *B. cereus sensu lato* (*s.l.*), is a species complex comprising numerous closely related lineages, which vary in their ability to cause illness in humans and animals. The classification of *B. cereus s.l.* isolates into species-level taxonomic units is essential for facilitating communication between and among microbiologists, clinicians, public health officials, and industry professionals, but is not always straightforward. A recently proposed genomospecies-subspecies-biovar taxonomic framework aims to provide a standardized nomenclature for this species complex but relies heavily on whole-genome sequencing (WGS), a technology with limited accessibility. It thus is unclear whether popular, low-cost typing methods (e.g., single- and multi-locus sequence typing) remain congruent with the proposed taxonomy. Here, we characterize 2,231 *B. cereus s.l.* genomes using a combination of *in silico* (i) average-nucleotide identity (ANI)-based genomospecies assignment, (ii) ANI-based subspecies assignment, (iii) seven-gene multi-locus sequence typing (MLST), (iv) *panC* group assignment, (v) *rpoB* allelic typing, and (vi) virulence factor detection. We show that sequence types (STs) assigned using MLST can be used for genomospecies assignment, and we provide a comprehensive list of ST/genomospecies associations. For *panC* group assignment, we show that an adjusted, eight-group framework is largely congruent with the proposed eight-genomospecies taxonomy and resolves incongruencies observed in the historical seven-group framework among isolates assigned to *panC* Groups II, III, and VI. We additionally provide a list of loci that capture the topology of the whole-genome *B. cereus s.l.* phylogeny that may be used in future sequence typing efforts. For researchers with access to WGS, MLST, and/or *panC* data, we showcase how our recently released software, BTyper3 (https://github.com/lmc297/BTyper3), can be used to assign *B. cereus s.l.* isolates to taxonomic units within this proposed framework with little-to-no user intervention or domain-specific knowledge of *B. cereus s.l.* taxonomy. We additionally outline a novel method for assigning *B. cereus s.l.* genomes to pseudo-gene flow units within proposed genomospecies. The results presented here highlight the backwards-compatibility and accessibility of the proposed taxonomic framework and illustrate that WGS is not a necessity for microbiologists who want to use the proposed taxonomy effectively.

## Introduction

The *Bacillus cereus* group, also known as *B. cereus sensu lato* (*s.l.*), is a species complex composed of numerous closely related, Gram-positive, spore-forming lineages with varying pathogenic potential (Rasko et al., 2005; Stenfors Arnesen et al., 2008; Messelhäußer and Ehling-Schulz, 2018; Ehling-Schulz et al., 2019). While some members of *B. cereus s.l.* have essential roles in agriculture and industry (e.g., as biocontrol agents) (Elshaghabee et al., 2017; Jouzani et al., 2017), others can cause illnesses with varying degrees of severity. Some members of the group, for example, are capable of causing severe forms of anthrax and anthrax-like illness that may result in death (Pilo and Frey, 2011; Moayeri et al., 2015; Pilo and Frey, 2018). Other members of the group can cause foodborne illness that manifests in either an emetic form (i.e., intoxication characterized by vomiting symptoms and an incubation period of 0.5 – 6 h) or diarrheal form (i.e., toxicoinfection characterized by diarrheal symptoms and an incubation period of 8 – 16 h) (Stenfors Arnesen et al., 2008; Ehling-Schulz et al., 2015; Messelhäußer and Ehling-Schulz, 2018; Rouzeau-Szynalski et al., 2020).

Differentiating beneficial *B. cereus s.l.* strains from those that are capable of causing illness or death thus requires microbiologists, clinicians, public health officials, and industry professionals to communicate the potential risk associated with a given isolate. However, the lack of a “common language” for describing *B. cereus s.l.* isolates has hindered communication between and among scientists and other professionals and could potentially lead to dangerous mischaracterizations of an isolate’s virulence potential. Anthrax-causing strains that possess phenotypic characteristics associated with “*B. cereus*” (e.g., motility, hemolysis on Sheep RBC) (Tallent et al., 2019), for example, have been referred to as “*B. anthracis*” (Leendertz et al., 2004), “*B. cereus*” (Hoffmaster et al., 2004; Avashia et al., 2007; Wilson et al., 2011), “*B. cereus* variety anthracis” (Klee et al., 2010), “*B. cereus* biovar anthracis” (Antonation et al., 2016; Marston et al., 2016), and “*B. cereus* biovar *anthracis*” or “*B. cereus* Biovar *anthracis*” (Brezillon et al., 2015; Antonation et al., 2016; Ehling-Schulz et al., 2019; Romero-Alvarez et al., 2020). Similarly, some *B. cereus s.l.* isolates that are closely related to emetic toxin (cereulide)-producing isolates are incapable of causing emetic intoxication themselves but can cause the diarrheal form of *B. cereus s.l.* illness (Ehling-Schulz et al., 2005; Jessberger et al., 2015; Riol et al., 2018; Carroll and Wiedmann, 2020). However, as there is no standardized name for these isolates, they have been referred to as “emetic-like *B. cereus*” (Ehling-Schulz et al., 2005), “*B. paranthracis*” (Liu et al., 2017; Bukharin et al., 2019), “*B. cereus*”, “Group III *B. cereus*” (i.e., assigned to Group III using the sequence of *panC* and the seven-phylogenetic group framework proposed by Guinebretiere, et al.), and “*B. cereus s.s.*” (although it should be noted that these strains do not fall within the genomospecies boundary of the *B. cereus s.s.* type strain and thus are not actually members of the *B. cereus s.s.* species) (Guinebretiere et al., 2010; Gdoura-Ben Amor et al., 2018; Glasset et al., 2018; Zhuang et al., 2019).

Recently, we proposed a standardized taxonomic nomenclature for *B. cereus s.l*. that is designed to minimize incongruencies and ambiguities within the *B. cereus s.l.* taxonomic space (Carroll et al., 2020). The proposed taxonomy consists of: (i) a standardized set of eight genomospecies names (i.e., *B. pseudomycoides, B. paramycoides, B. mosaicus, B. cereus s.s., B. toyonensis, B. mycoides, B. cytotoxicus*) that correspond to resolvable, non-overlapping genomospecies clusters obtained at a ≈92.5 average nucleotide identity (ANI) breakpoint; (ii) a formal collection of two subspecies names which account for established lineages of medical importance (i.e., subspecies *anthracis*, which is used to refer to the classic non-motile, non-hemolytic lineage referred to as “*B. anthracis*”, and subspecies *cereus*, which is used to refer to *panC* Group III lineages that encompass cereulide-producing isolates [i.e., “emetic *B. cereus*”] and the non-cereulide-producing isolates interspersed among them); and (iii) a standardized collection of biovar terms (i.e., Anthracis, Emeticus, Thuringiensis), which can be used to account for the heterogeneity of clinically and industrially important phenotypes (i.e., production of anthrax toxin, cereulide, and/or insecticidal crystal proteins, respectively) (Carroll et al., 2020). However, this nomenclatural framework was developed using data derived from whole-genome sequencing (WGS) efforts, a technology that may not be accessible to all microbiologists or necessary for all microbiological studies. Hence, an assessment of congruency between WGS-based and single- or multi-locus sequencing-based genotyping and taxonomic assignment methods is needed. Here, we characterize 2,231 *B. cereus s.l.* genomes using a combination of *in silico* (i) ANI-based genomospecies assignment, (ii) ANI-based subspecies assignment, (iii) seven-gene multi-locus sequence typing (MLST), (iv) *panC* group assignment, (v) *rpoB* allelic typing, and (vi) virulence factor detection to show that popular, low-cost typing methods (e.g., single- and MLST) remain largely congruent with the proposed taxonomy. We additionally showcase how our recently released software, BTyper3 (Carroll et al., 2020), can be used to assign *B. cereus s.l.* isolates to taxonomic units within this proposed framework using WGS, MLST, and/or *panC* data. Further, we provide a list of loci that mirror the topology of the whole-genome *B. cereus s.l.* phylogeny, which may be used in future sequence typing efforts. Finally, we provide a novel method for assigning *B. cereus s.l.* isolates to pseudo-gene flow units using WGS data. The results presented here showcase that the proposed taxonomic framework for *B. cereus s.l.* is backwards-compatible with historical *B. cereus s.l.* typing efforts and can be utilized effectively, regardless of whether WGS is used to characterize isolates or not.

## Methods

### Acquisition of *Bacillus cereus s.l.* genomes

All genomes submitted to the National Center for Biotechnology Information (NCBI) RefSeq (Pruitt et al., 2007) database as a published *B. cereus s.l.* species (Lechner et al., 1998; Guinebretiere et al., 2013; Jimenez et al., 2013; Miller et al., 2016; Liu et al., 2017; Carroll et al., 2020) were downloaded (*n* = 2,231, accessed November 19, 2018; Supplementary Table S1). QUAST v. 5.0.2 (Gurevich et al., 2013) and CheckM v. 1.0.7 (Parks et al., 2015) were used to assess the quality of each genome, and BTyper3 v. 3.1.0 was used to assign each genome a genomospecies, subspecies (if applicable), and biovar(s) (if applicable), using a recently proposed taxonomy (Carroll et al., 2020). Genomes with (i) N50 > 100,000, (ii) CheckM completeness ≥ 97.5%, (iii) CheckM contamination ≤ 2.5%, and (iv) a genomospecies assignment that corresponded to a published *B. cereus s.l.* genomospecies were used in subsequent steps unless otherwise indicated (Supplementary Table S1). Genomes that did not meet these quality thresholds, as well as those which were assigned to an unknown or unpublished genomospecies (i.e., “Unknown *B. cereus* group Species 13-18” described previously) (Carroll et al., 2020) or an effective or proposed *B. cereus s.l.* genomospecies (i.e., “*B. bingmayongensis*”, “*B. clarus*”, *“B. gaemokensis*”, or *“B. manliponensis*”), were excluded (Jung et al., 2010; Jung et al., 2011; Liu et al., 2014; Acevedo et al., 2019), yielding a set of 1,741 high-quality *B. cereus s.l.* genomes. All subsequent analyses relied on one of two sets of genomes, as indicated: (i) the full set of 2,231 *B. cereus s.l.* RefSeq genomes, or (ii) the set of 1,741 high-quality genomes, with effective, proposed, unknown, and unpublished genomospecies removed. In some cases, the type strain genome of effective *B. cereus s.l.* species “*B. manliponensis*” was used to root a phylogeny, as it is the most distantly related member of the species complex (Jung et al., 2011; Carroll et al., 2020).

### Average nucleotide identity calculations, genomospecies cluster delineation, and identification of medoid genomes

FastANI v. 1.0 (Jain et al., 2018) was used to calculate pairwise ANI values between all 1,741 high-quality *B. cereus s.l.* genomes (see section “Acquisition of *Bacillus cereus s.l.* genomes” above). Genomospecies clusters and their respective medoid genomes were identified among all 1,741 genomes at all previously proposed ANI genomospecies thresholds for *B. cereus s.l.* (i.e., thresholds of 92.5, 94, 95, and 96 ANI) (Guinebretiere et al., 2013; Jimenez et al., 2013; Miller et al., 2016; Liu et al., 2017; Carroll et al., 2020) as described previously (Carroll et al., 2020), using the bactaxR package (Carroll et al., 2020) in R v. 3.6.1 (R Core Team, 2019) and following dependencies: ape v. 5.3 (Paradis et al., 2004; Paradis and Schliep, 2019), cluster v. 2.1.0 (Maechler et al., 2019), dendextend v. 1.13.4 (Galili, 2015), dplyr v. 0.8.5 (Wickham et al., 2020), ggplot2 v. 3.3.0 (Wickham, 2016), ggtree v. 1.16.6 (Yu et al., 2017; Yu et al., 2018), igraph v. 1.2.5 (Csardi and Nepusz, 2006), phylobase v. 0.8.10 (R Hackathon, 2019), phytools v. 0.7-20 (Revell, 2012), readxl v. 1.3.1 (Wickham and Bryan, 2019), reshape2 v. 1.4.4 (Wickham, 2007), and viridis v. 0.5.1 (Garnier, 2018).

FastANI was additionally used to calculate ANI values between each of the 2,231 genomes in the full set of *B. cereus s.l.* genomes and the type strain genomes of all 21 published and effective *B. cereus s.l.* species described prior to 2020 (Supplementary Table S1) so that the historical practice of assigning *B. cereus s.l.* genomes to species using type strain genomes could be assessed.

### Construction of *B. cereus s.l.* whole-genome phylogeny

To remove highly similar genomes and reduce the full set of 1,741 high-quality genomes to a smaller set of genomes that encompassed the diversity of *B. cereus s.l.* in its entirety, medoid genomes were identified among the set of 1,741 high-quality *B. cereus s.l.* genomes (see section “Acquisition of *Bacillus cereus s.l.* genomes” above) at a 99 ANI threshold using the bactaxR package in R (see section “Average nucleotide identity calculations, genomospecies cluster delineation, and identification of medoid genomes” above). Core single-nucleotide polymorphisms (SNPs) were identified among the resulting set of non-redundant genomes (*n* = 313; Supplementary Table S1) using kSNP3 v. 3.92 (Gardner and Hall, 2013; Gardner et al., 2015) and the optimal *k*-mer size reported by Kchooser (*k* = 19). IQ-TREE v. 1.5.4 (Nguyen et al., 2015) was used to construct a maximum likelihood phylogeny using the resulting core SNPs, the General Time-Reversible (Tavaré, 1986) nucleotide substitution model with a gamma rate-heterogeneity parameter (Yang, 1994) and ascertainment bias correction (Lewis, 2001) (i.e., the GTR+G+ASC nucleotide substitution model), and 1,000 replicates of the ultrafast bootstrap approximation (Minh et al., 2013; Hoang et al., 2018). The aforementioned core SNP detection and phylogeny construction steps were then repeated among the same set of 313 medoid genomes, with the addition of the “*B. manliponensis*” type strain genome (*n* = 314). The resulting phylogenies were annotated using the bactaxR package in R.

### Construction of *panC, rpoB*, and seven-gene MLST phylogenies

BTyper v. 2.3.2 (Carroll et al., 2017) was used to extract the nucleotide sequences of (i) *panC*, (ii) *rpoB*, and (iii) the seven genes used in the PubMLST (Jolley and Maiden, 2010; Jolley et al., 2018) MLST scheme for *B. cereus* (i.e., *glp, gmk, ilv, pta, pur, pyc*, and *tpi*) from each of the 1,741 high-quality *B. cereus s.l.* genomes. MAFFT v. 7.453-with-extensions (Katoh et al., 2002; Katoh and Standley, 2013) was used to construct an alignment for each gene, and IQ-TREE was used to build a ML phylogeny from each resulting alignment, as well as an alignment constructed by concatenating the seven MLST genes, using the optimal nucleotide substitution model selected using ModelFinder (Kalyaanamoorthy et al., 2017) and 1,000 replicates of the ultrafast bootstrap approximation. The resulting phylogenies were annotated using the bactaxR package in R.

### Construction of the adjusted, eight-group *panC* group assignment framework

Medoid genomes were identified among the full set of 1,741 high-quality *B. cereus s.l.* genomes at a 99 ANI threshold (*n* = 313; see section “Average nucleotide identity calculations, genomospecies cluster delineation, and identification of medoid genomes” above). BTyper v. 2.3.3 was used to extract *panC* from each of the 313 *B. cereus s.l.* genomes, and MAFFT v. 7.453-with-extensions was used to construct an alignment. RhierBAPS v. 1.1.3 (Tonkin-Hill et al., 2018) was used to identify *panC* clusters within the alignment using two clustering levels; the nine top level (i.e., Level 1) clusters were used in subsequent steps, as they most closely mirrored the original seven *panC* groups (24 separate clusters were produced at Level 2). BTyper v. 2.3.3 was then used to extract *panC* from the full set of high-quality *B. cereus s.l.* genomes (*n* = 1,741; note that *panC* could not be extracted from all genomes), and the cd-hit-est command from CD-HIT v. 4.8.1 (Li and Godzik, 2006; Fu et al., 2012) was then used to cluster the resulting *panC* genes at a sequence identity threshold of 0.99. *panC* sequences that fell within the same CD-HIT cluster as a *panC* sequence from one or more of the 313 medoid genomes (*n* = 1,736) were assigned the RhierBAPS cluster of the medoid genome(s). MAFFT was used to construct an alignment of all 1,736 *panC* genes, and IQ-TREE v. 1.6.5 was used to construct a phylogeny using the resulting alignment as input, the optimal nucleotide substitution model selected using ModelFinder (i.e., the TVM+F+R4 model), and 1,000 replicates of the ultrafast bootstrap approximation.

The nine Level 1 RhierBAPS *panC* cluster assignments were then manually compared to *panC* groups assigned using BTyper v. 2.3.3 and the legacy seven-group framework. RhierBAPS *panC* groups were then re-named so that they most closely resembled the historical group assignments used by Guinebretiere, et al. and BTyper v. 2.3.3 (Guinebretiere et al., 2008; Guinebretiere et al., 2010; Carroll et al., 2017).

### Identification of putative loci for future single- and MLST efforts

Prokka v. 1.12 (Seemann, 2014) was used to annotate each of the 313 *B. cereus s.l.* medoid genomes identified at 99 ANI (see section “Average nucleotide identity calculations, genomospecies cluster delineation, and identification of medoid genomes” above), and the resulting protein sequences were divided randomly into 11 sets (ten sets containing 30 genomes, and one set containing 13 [the remainder] genomes) (Carroll et al., 2020). OrthoFinder v. 2.3.3 (Emms and Kelly, 2015) was used to identify single-copy core genes present among all genomes in each set, and, subsequently, among all 313 genomes, using the iterative approach described previously (Carroll et al., 2020). Nucleotide sequences of each of the 1,719 single-copy core genes present among all 313 genomes were aligned using MAFFT v. 7.453-with-extensions, and each resulting gene alignment was used as input for IQ-TREE v. 1.6.5. A maximum likelihood phylogeny was constructed for each gene using the GTR+G nucleotide substitution model and 1,000 replicates of the ultrafast bootstrap approximation.

The Kendall-Colijn (Kendall and Colijn, 2015; 2016; Jombart et al., 2017) test described by Katz, et al. (Katz et al., 2017) was used to assess the topological congruency between phylogenies constructed using each core gene and the “true” *B. cereus s.l.* whole-genome phylogeny (see section “Construction of *B. cereus s.l.* whole-genome phylogeny” above). For each topological comparison, both phylogenies were rooted at the midpoint, and a lambda value of 0 (to give weight to tree topology rather than branch lengths) (Katz et al., 2017) and background distribution of 1,000 random trees were used. A phylogeny was considered to be more topologically similar to the “true” *B. cereus s.l.* whole-genome phylogeny than would be expected by chance if a significant *P*-value (*P* < 0.05) resulted after a Bonferroni correction was applied (Katz et al., 2017).

Metrics used for assessing the quality of putative typing loci included (i) length of the longest, uninterrupted/ungapped stretch of continuous sequence within the gene alignment, (ii) proportion of sites within the gene alignment that did not include gaps, (iii) proportion of the gene alignment that was covered by the longest uninterrupted/ungapped stretch of continuous sequence, and (iv) Bonferroni-corrected Kendall-Colijn *P*-value (Supplementary Table S2). Each individual gene was then detected within the full set of 1,741 high-quality *B. cereus s.l.* genomes (see section “Acquisition of *Bacillus cereus s.l.* genomes” above) using nucleotide BLAST v. 2.9.0 (Camacho et al., 2009), as implemented in BTyper v. 2.3.3, by aligning the alleles of each single-copy core gene (*n* = 313) to each of the 1,741 genomes. A final set of candidate loci for single- and MLST was then identified (*n* = 255). A gene was included in the final set if: (i) ≥ 90% of the sites within the gene’s alignment did not contain gap characters; (ii) the longest stretch of uninterrupted/ungapped continuous sequence within the gene’s alignment covered ≥ 90% of the full length of the gene’s alignment; (iii) the maximum likelihood phylogeny constructed using the gene as input was topologically similar to the “true” whole-genome phylogeny (i.e., Kendall-Colijn *P*-value < 0.05 after a Bonferroni correction); (iv) a single copy of the gene could be detected in all 1,741 high-quality *B. cereus s.l.* genomes, using minimum percent nucleotide identity and coverage thresholds of 90% each and a maximum E-value threshold of 1E-5 (Supplementary Table S2).

### Functional annotation of putative loci for future single- and MLST efforts

Amino acid sequences of the resulting 255 candidate loci (see section “Identification of putative loci for future single- and MLST efforts”; Supplementary Table S2) were functionally annotated using eggNOG mapper v. 2.0 (Huerta-Cepas et al., 2017; Huerta-Cepas et al., 2019). The resulting Clusters of Orthologous Groups (COG) functional categories were visualized in R using the igraph v. 1.2.5 package (Csardi and Nepusz, 2006). The GOGO Webserver (http://dna.cs.miami.edu/GOGO/; accessed May 30, 2020) was used to calculate pairwise semantic/functional similarities between genes based on their assigned Gene Ontology (GO) terms and to cluster genes based on their GO term similarities (Zhao and Wang, 2018). For each of the three GO directed acyclic graphs (i.e., Biological Process Ontology, Cellular Component Ontology, and Molecular Function Ontology) (Ashburner et al., 2000; The Gene Ontology Consortium, 2018), an *n* × *n* matrix of pairwise similarities produced by GOGO were converted into a dissimilarity matrix by subtracting all values from an *n* × *n* matrix containing 1.0s. Non-metric multidimensional scaling (NMDS) (Kruskal, 1964) was performed using the resulting dissimilarity matrix, the metaMDS function in the vegan (Oksanen et al., 2019) package in R, two dimensions (*k* = 2), and a maximum of 10,000 random starts. Convergent solutions were reached in under 100 random starts for the biological process and cellular component dissimilarity matrices and in under 1,400 random starts for the molecular function dissimilarity matrix. The results from each NMDS run were plotted in R using ggplot2.

### Identification of microbial gene flow units using recent gene flow and implementation of the pseudo-gene flow unit assignment method in BTyper3 v. 3.1.0

The “PopCOGenT” module available in PopCOGenT (downloaded October 5, 2019) (Arevalo et al., 2019) was used to identify gene flow units (i.e., “main clusters” reported by PopCOGenT) among the 313 *B. cereus s.l.* medoid genomes identified at 99 ANI (Figure 1A; see section “Average nucleotide identity calculations, genomospecies cluster delineation, and identification of medoid genomes” above), using the following dependencies: Mugsy v. v1r2.3 (Angiuoli and Salzberg, 2011) and Infomap v. 0.2.0 (Rosvall et al., 2009).

**Figure 1.**
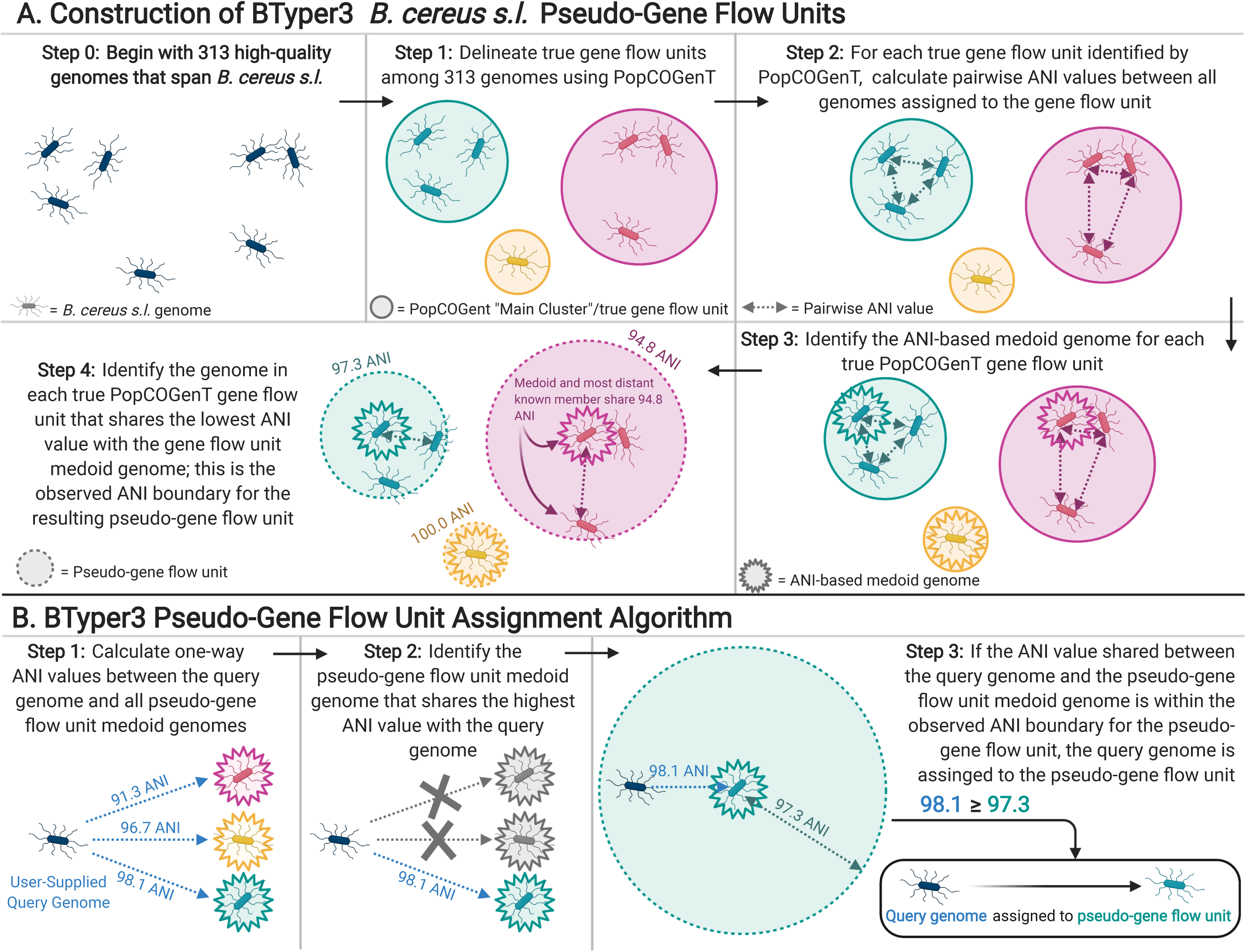
(A) Graphical depiction of the methods used to construct *B. cereus s.l.* pseudo-gene flow units used by BTyper3. The 313 high-quality *B. cereus s.l.* genomes (Step 0) were medoid genomes identified among a set of 1,741 high-quality *B. cereus s.l.* genomes at a 99 average nucleotide identity (ANI) threshold using the bactaxR package in R. This step was performed to remove highly similar genomes and reduce the full set of 1,741 high-quality genomes to a smaller set of genomes that encompassed the diversity of *B. cereus s.l.* in its entirety. Gene flow units delineated using PopCOGenT (Step 1) were the “main clusters” reported by the PopCOGenT module. (B) Graphical depiction of the pseudo-gene flow unit assignment algorithm implemented in BTyper3. Pseudo-gene flow unit medoid genomes (Steps 1-3) are the output of the steps outlined in (A). If a user-supplied query genome does not fall within the observed ANI boundary of the most similar pseudo-gene flow unit medoid genome (Step 3), the second-through-fifth most similar pseudo-gene flow unit medoid genomes are queried. All ANI values were calculated using FastANI. The figure was created with BioRender (https://biorender.com/).

Pairwise ANI values were then calculated between genomes within each of the 33 PopCOGenT gene flow units using FastANI v. 1.0, and bactaxR was used to identify the medoid genome for each PopCOGenT gene flow unit based on the resulting pairwise ANI values (Figure 1A). The minimum ANI value shared between the PopCOGenT gene flow unit medoid genome and all other genomes assigned to the same gene flow unit using PopCOGenT was treated as the observed ANI boundary for the gene flow unit; the observed ANI boundary formed by a PopCOGenT gene flow unit medoid genome forms what we refer to here as a pseudo-gene flow unit (Figure 1A).

The 33 resulting medoid genomes for each of the 33 pseudo-gene flow units, as well as the genomes of effective and proposed *B. cereus s.l.* species, were then used to create a rapid pseudo-gene flow unit typing scheme in BTyper3 v. 3.1.0 (Figure 1). For this approach, ANI values are calculated between a user’s query genome and the set of 33 pseudo-gene flow unit medoid genomes using FastANI (Figure 1B and Figure 2). The closest-matching medoid genome and its ANI value relative to the query are identified; additionally, the previously observed ANI boundaries for the medoid genome’s respective pseudo-gene flow unit are reported (Figure 1B and Figure 2). It is important to note that this pseudo-gene flow unit assignment method measures genomic similarity via ANI, which is fundamentally and conceptually very different from the methods that PopCOgenT employs. The ANI-based pseudo-gene flow unit assignment method described here does not query recent gene flow, nor does it use PopCOgenT or the methods that it employs directly. Thus, it cannot directly assign a genome to a PopCOgenT gene flow unit, and results should not be interpreted as a true measurement of gene flow. However, this approach allows researchers to rapidly identify the closest medoid genome of previously delineated true gene flow units (Figure 1A), based on a metric of genomic similarity, which provides insight into the phylogenomic placement of a query genome within a larger *B. cereus s.l.* genomospecies.

**Figure 2.**
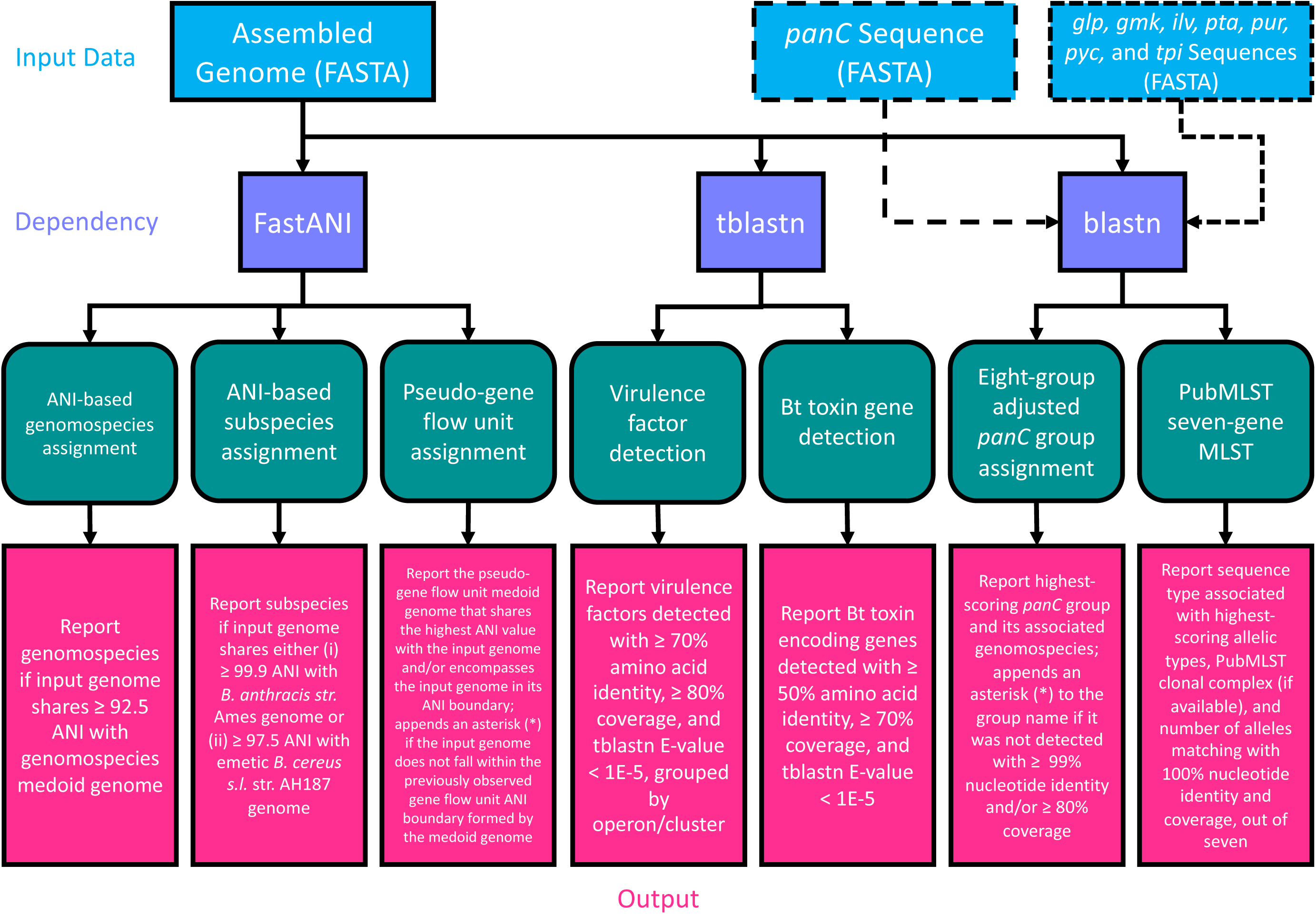
Flow chart describing the workflow implemented in BTyper3 v. 3.1.0. Input data in FASTA format (blue boxes) can consist of any of the following: (i) a whole genome (complete or draft; can be used with all/all combinations of workflow steps), (ii) a *panC* sequence (can be used with the eight-group adjusted *panC* group assignment workflow only), or (iii) sequences of the seven loci used in PubMLST’s seven-gene multi-locus sequence typing (MLST) scheme for *B. cereus s.l.* (sequences can be in multi-FASTA format, or concatenated into a single sequence; can be used with the PubMLST seven-gene MLST workflow only). Purple boxes represent software dependencies required for each type of input data, while green boxes represent the various analyses that can be conducted in BTyper3 v. 3.1.0. Pink boxes denote the output that BTyper3 v. 3.1.0 reports for each analysis.

### Implementation of virulence factor detection in BTyper3 v. 3.1.0

Versions of BTyper3 prior to v. 3.1.0 (Carroll et al., 2020), as well as the original BTyper (i.e., BTyper v. 2.3.3 and earlier) (Carroll et al., 2017) detected virulence factors using translated nucleotide BLAST (Camacho et al., 2009) and minimum amino acid identity and coverage thresholds of 50% and 70%, respectively, as these values had been shown to correlate with PCR-based detection of virulence factors (Kovac et al., 2016). However, these thresholds were selected using a limited number of *B. cereus s.l.* isolates with limited genomic diversity and can potentially lead to the detection of remote homologs that do not correlate with a virulence phenotype (i.e., false positive hits). For example, some *B. cereus s.l.* isolates possess a gene that shares a low degree of homology with *cesC*, but still meet these virulence factor detection thresholds (see Figure 5 of Carroll, et al., 2017) (Carroll et al., 2017). Users with limited knowledge of *B. cereus s.l.* virulence factors, or those who do not know how to interpret BLAST identity and coverage thresholds, may infer that these isolates have a potential to produce cereulide, when they actually do not. A similar phenomenon is observed with some members of the “*B. cereus*” exo-polysaccharide capsule (Bps)-encoding genes (e.g., *bpsEF*) (Carroll et al., 2019).

**Figure 3.**
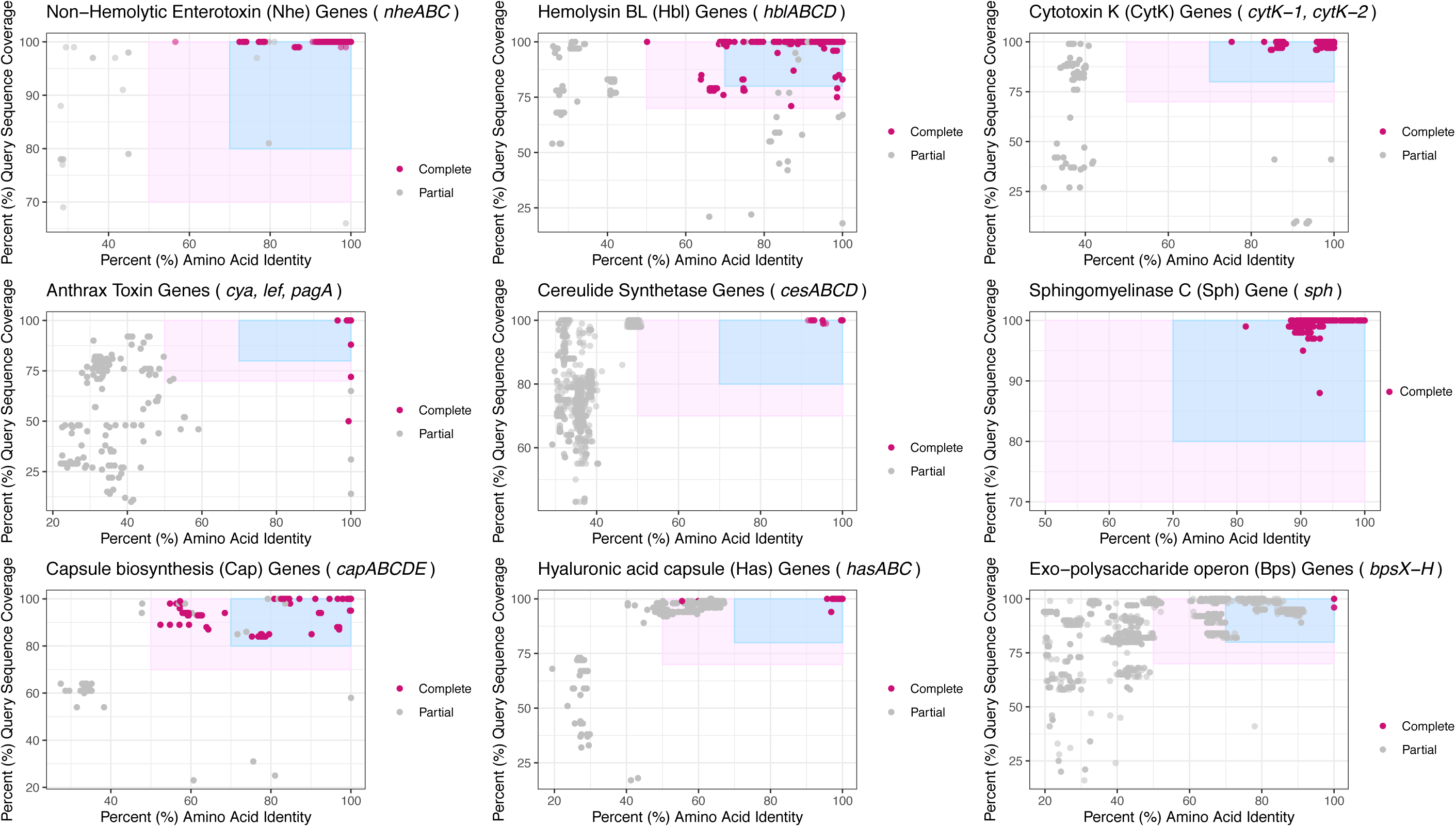
Virulence factors detected in 1,741 high-quality *B. cereus s.l.* genomes at various minimum percent amino acid identity (X-axes) and query sequence coverage (Y-axes) thresholds. Each subplot denotes a virulence factor composed of one or more genes listed in the subplot title. Points represent the individual genes listed in the subplot title. The light pink rectangle denotes amino acid identity and coverage values at which the original BTyper (Btyper v. 2.3.3 and previous versions) would report a gene as “present” (i.e., 50 and 70% amino acid identity and coverage thresholds, respectively). The blue rectangle denotes the updated virulence factor cutoffs used by BTyper3 v. 3.1.0 (i.e., 70 and 80% amino acid identity and coverage thresholds, respectively). Points shaded in dark pink (i.e., “Complete”) were (i) detected within a *B. cereus s.l.* genome at the default minimum amino acid identity and coverage thresholds used by the original BTyper (i.e., BTyper v. 2.3.3, at 50 and 70%, respectively), and (ii) were part of a “complete” virulence factor, as listed in the subplot title (i.e., all other genes comprising the virulence factor were detected in the genome at the 50 and 70% minimum amino acid identity and coverage thresholds used by BTyper v. 2.3.3, respectively). Points colored in gray (i.e., “Partial”) denote genes that were not detected at the 50 and 70% minimum amino acid identity and coverage thresholds used by BTyper v. 2.3.3 and/or were part of a virulence factor that was not present in its entirety in the respective genome at 50 and 70% amino acid identity and coverage, respectively. All genes were detected using BTyper3 v. 3.1.0 with a minimum E-value threshold of 1E-5.

**Figure 4.**
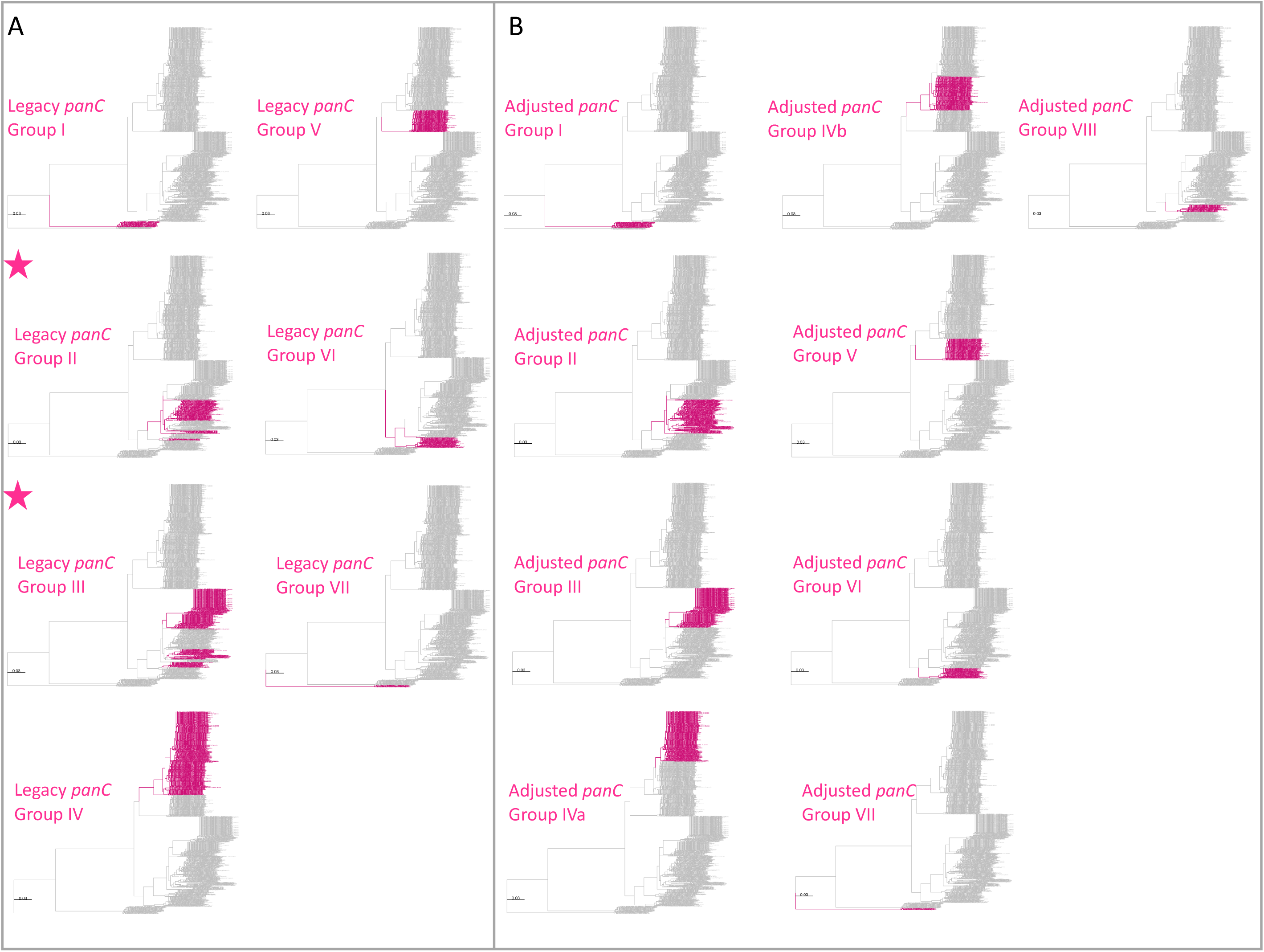
Maximum likelihood phylogeny constructed using *panC*, extracted from 1,736 high-quality *B. cereus s.l.* genomes. Branches and tip labels are colored by (A) *panC* group (I-VII), assigned using the *panC* group assignment method implemented in the original BTyper v. 2.3.3 (i.e., the “legacy” *panC* group assignment method), and (B) adjusted *panC* group assignment (I-VIII), obtained using RhierBAPS Level 1 cluster assignments for *panC*. For all *panC* group assignments, the “foreground” *panC* group is colored (pink) and the background *panC* groups are shown in gray. Phylogenies for which the foreground *panC* group (pink) presents as polyphyletic are annotated with a pink star in the upper left corner of the panel. Phylogenies are rooted using the *panC* sequence of the *“B. manliponensis*” type strain (omitted for clarity), and branch lengths are reported in substitutions per site. IQ-TREE v. 1.6.5 was used to construct a phylogeny, using the optimal nucleotide substitution model selected using ModelFinder (i.e., the TVM+F+R4 model).

**Figure 5.**
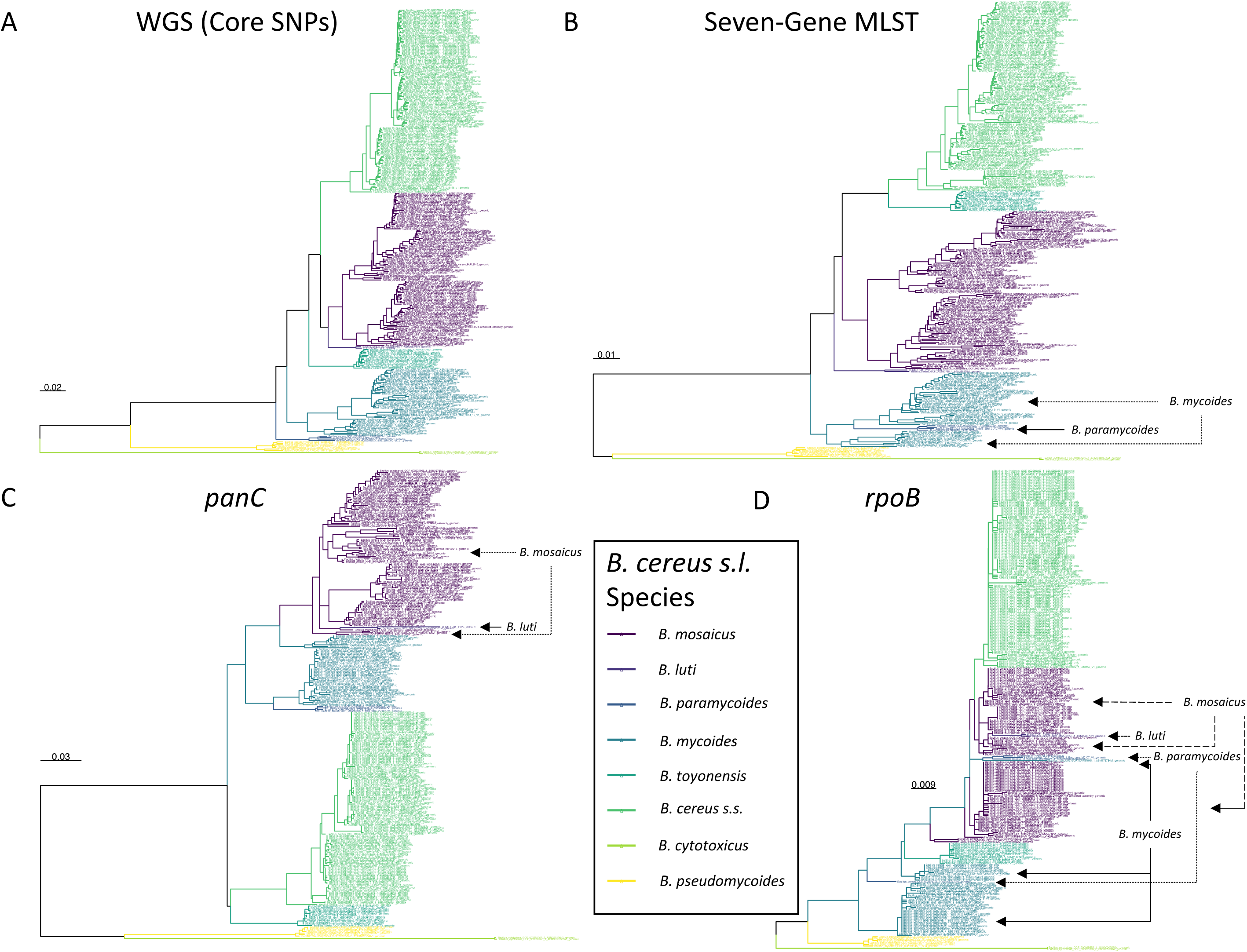
Maximum likelihood phylogenies constructed using (A) genome-wide core SNPs (WGS), (B) seven concatenated multi-locus sequence typing (MLST) genes, (C) *panC*, and (D) *rpoB* identified among 313 high-quality *B. cereus s.l.* medoid genomes identified at a 99 average nucleotide identity (ANI) threshold. Branches and tip labels are colored by ANI-based genomospecies assignment using the proposed *B. cereus s.l.* taxonomic framework (i.e., eight genomospecies assigned using medoid genomes and a 92.5 ANI threshold). Polyphyletic genomospecies and the genomospecies interspersed among them are annotated with arrows. Phylogenies are rooted at the midpoint, and branch lengths are reported in substitutions per site. Genomospecies were assigned using BTyper3 v. 3.1.0 and FastANI v. 1.0. For the WGS tree (A), core SNPs were identified among all 313 *B. cereus s.l.* genomes using kSNP3 v. 3.92 and the optimal *k*-mer size reported by Kchooser (*k* = 19). For the MLST, *panC*, and *rpoB* trees (B,C, and D, respectively), BTyper v. 2.3.3 was used to extract all loci from the set of 313 *B. cereus s.l.* genomes, and MAFFT v. 7.453-with-extensions was used to construct an alignment for each locus. For each alignment, IQ-TREE v. 1.5.4 (A) and 1.6.5 (B, C, and D) was used to construct a phylogeny, using either the GTR+G+ASC nucleotide substitution model (A), or the optimal model selected using ModelFinder (B, C, and D).

To improve *in silico* virulence factor detection in *B. cereus s.l.* genomes, the BTyper3 v. 3.1.0 virulence factor database was constructed to include amino acid sequences of the following virulence factors: (i) anthrax toxin genes *cya, lef*, and *pagA* (the same sequences used for assignment of biovar Anthracis in all previous versions of BTyper3); (ii) cereulide synthetase genes *cesABCD* (the same sequences used for assignment of biovar Emeticus in all previous versions of BTyper3); (iii) non-hemolytic enterotoxin (Nhe) genes *nheABC* (used in the original BTyper v. 2.3.3 and earlier); (iv) hemolysin BL (Hbl) genes *hblABCD* (used in the original BTyper v. 2.3.3 and earlier); (v) cytotoxin K (CytK) variant 1 and 2 (*cytK-1* and *cytK-2*, respectively; used in the original BTyper v. 2.3.3 and earlier); (vi) sphingomyelinase Sph gene *sph* (used in the original BTyper v. 2.3.0-2.3.3); (vii) anthrax capsule biosynthesis (Cap) genes *capABCDE* (used in the original BTyper v. 2.3.3 and earlier); (viii) hyaluronic acid capsule (Has) genes *has ABC* (*hasA* was included in the original BTyper v. 2.3.3 and earlier, and *hasBC* were added here) (Oh et al., 2011); (ix) exo-polysaccharide capsule (Bps) genes *bpsXABCDEFGH* (used in the original BTyper v 2.0.1-2.3.3).

To provide updated boundaries for virulence factor detection based on a larger set of genomes that span *B. cereus s.l.*, BTyper3 v. 3.1.0 was used to identify all virulence factors listed above in the complete set of 1,741 high-quality genomes (see section “Acquisition of *Bacillus cereus s.l.* genomes” above), using a maximum BLAST E-value threshold of 1E-5 (Carroll et al., 2017; Carroll et al., 2020), but with minimum amino acid identity and coverage thresholds of 0% each. Plots of virulence factors detected within all genomes at various amino acid identity and coverage thresholds were constructed using ggplot2 in R (Figure 3 and Supplementary Table S3). Based on these plots, amino acid identity and coverage thresholds of 70% and 80%, respectively, were implemented as the default thresholds for virulence factor detection in BTyper3 v. 3.1.0 (Figure 3). Additionally, to reduce the risk of users mis-interpreting spurious hits that do not correlate with a virulence phenotype, BTyper3 v. 3.1.0 reports virulence factors at an operon/cluster level; for example, if only *cesC* is detected in a genome, BTyper3 reports that only one of four cereulide synthetase-encoding genes were detected (Figure 2). Similarly, some *B. cereus s.l.* isolates possess genes that share a high degree of homology with Bps-encoding genes (e.g., > 90% identity and coverage); to avoid users mis-interpreting that this isolate may produce a Bps capsule, BTyper3 reports the fraction of *bps* hits out of nine *bps* genes (Figure 2).

### Implementation of seven-gene MLST in BTyper3 v. 3.1.0

The PubMLST seven-gene MLST scheme for *B. cereus s.l.* implemented in the original version of BTyper (i.e., BTyper v. 2.3.3 and earlier) was implemented in BTyper3 v. 3.1.0 as described previously (Carroll et al., 2017). The option to download the latest version of the *B. cereus s.l.* MLST database from PubMLST was also included in BTyper3 v. 3.1.0. Additionally, the clonal complex associated with each sequence type listed in PubMLST (if available), as well as the number of alleles that matched an allele in the PubMLST database with 100% identity and coverage out of seven, is reported in the BTyper3 final report (Figure 2).

### Implementation of *panC* group assignment in BTyper3 v. 3.1.0

The updated eight-group *panC* group assignment framework developed here (see section “Construction of the adjusted, eight-group *panC* group assignment framework” above) was used to construct a typing method in BTyper3 v. 3.1.0 (Figures 2 and 4). Briefly, BTyper3 v. 3.1.0 assigns a genome to a *panC* group using a database of 64 representative *panC* sequences from the 1,736 *B. cereus s.l. panC* sequences clustered at a 99% identity threshold described above. *panC* sequences of effective and proposed *B. cereus s.l.* species are also included in the database but are assigned a species name (e.g., “Group_manliponensis”) rather than a number (i.e., Group_I to Group_VIII). Nucleotide BLAST is used to align a query genome to the *panC* database, and the *panC* group producing the highest BLAST bit score is reported. Species associated with each *panC* group within the eight-group framework are also reported: (i) Group I (*B. pseudomycoides*), (ii) Group II (*B. mosaicus/B. luti*); (iii) Group III (*B. mosaicus*); (iv) Group IV (*B. cereus s.s.*); (v) Group V (*B. toyonensis*); (vi) Group VI (*B. mycoides/B. paramycoides*); (vii) Group VII (*B. cytotoxicus*); (viii) Group VIII (*B. mycoides*; Figure 2). If a query genome does not share ≥ 99% nucleotide identity and/or ≥ 80% coverage with one or more *panC* alleles in the database, the closest-matching *panC* group is reported with an asterisk (*).

### BTyper3 code availability

BTyper3, its source code, and its associated databases are free and publicly available at https://github.com/lmc297/BTyper3.

## Results

### Genomospecies defined using historical ANI-based genomospecies thresholds and species type strains are each integrated into one of eight proposed *B. cereus s.l.* genomospecies

Genomospecies assigned using higher, historical species cutoffs (i.e., 94, 95, and 96 ANI) and the type strain genomes of the 18 published *B. cereus s.l.* species described prior to 2020 were safely integrated into proposed genomospecies delineated at 92.5 ANI without polyphyly (Supplementary Table S4). Five of the eight genomospecies (i.e., *B. pseudomycoides, B. paramycoides, B. toyonensis, B. cytotoxicus*, and *B. luti*) encompassed all genomes assigned to the respective species using its type strain (Table 1), regardless of whether a 94, 95, or 96 ANI threshold was used. The remaining three genomospecies (i.e., *B. mosaicus, B. cereus s.s.*, and *B. mycoides*) simply integrated multiple species assigned using historical ANI-based genomospecies thresholds into a single genomospecies (Table 1). Regardless of whether a threshold of 94, 95, or 96 ANI was used, all genomes assigned to any of *B. albus, anthracis, mobilis, pacificus, paranthracis, tropicus*, and *wiedmannii* using species type strain genomes belonged to *B. mosaicus* (Table 1 and Supplementary Table S4). Likewise, all genomes assigned to any of *B. mycoides, nitratireducens, proteolyticus*, and *weihenstephanensis* using species type strain genomes and genomospecies thresholds of 94-96 ANI were assigned to the *B. mycoides* genomospecies cluster (Table 1 and Supplementary Table S4). Additionally, all genomes that shared 94-96 ANI with the *B. cereus s.s.* str. ATCC 14579 and/or *B. thuringiensis* serovar berliner str. ATCC 10792 type strain genomes belonged to the *B. cereus s.s.* genomospecies cluster (Table 1 and Supplementary Table S4). However, it should be noted that the “*B. cereus*” and “*B. thuringiensis*” species as historically defined are polyphyletic, and other strains often referred to as “*B. cereus*” or “*B. thuringiensis*” belong to other genomospecies clusters; emetic reference strain “*B. cereus*” str. AH187, for example, belongs to *B. mosaicus* and not *B. cereus s.s.* (Carroll et al., 2019; Carroll et al., 2020).

**Table 1.**
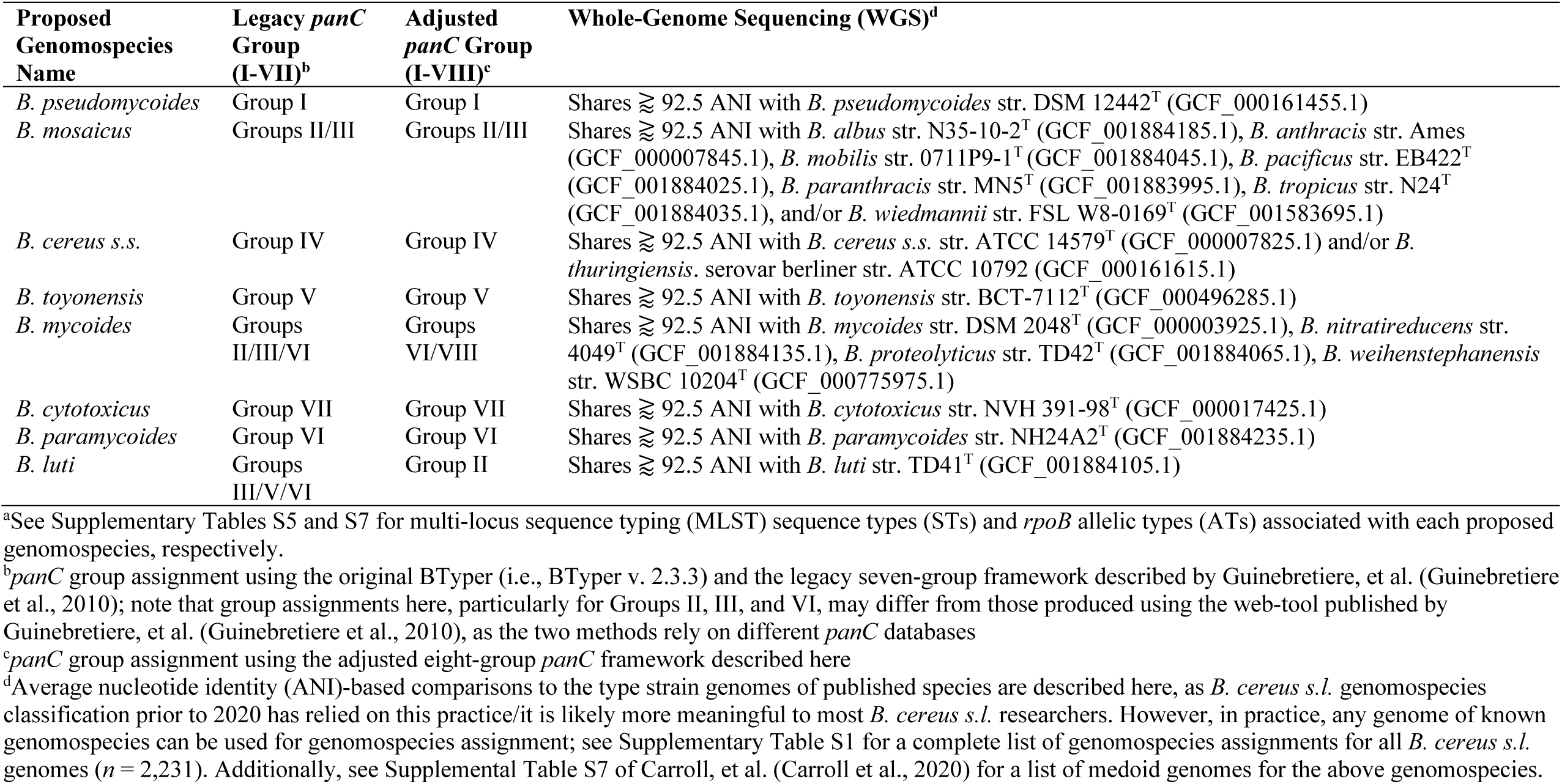
Proposed genomospecies-level taxonomy for *B. cereus s.l.* isolates.^a^

### STs assigned using seven-gene MLST can be used for *B. cereus s.l.* genomospecies assignment

All STs assigned using BTyper3 and PubMLST’s seven-gene MLST scheme *for B. cereus s.l*. (Jolley and Maiden, 2010; Jolley et al., 2018) were contained within a single proposed *B. cereus s.l.* genomospecies, and no STs were split across multiple genomospecies (Supplementary Tables S1 and S5). As such, a comprehensive list of ST/genomospecies associations for all NCBI RefSeq *B. cereus s.l.* genomes is available (*n* = 2,231; RefSeq accessed November 19, 2018, PubMLST *B. cereus* database accessed April 26, 2020; Supplementary Tables S1 and S5). However, it is essential to note that the *B. cereus s.l.* phylogeny constructed using the sequences of these seven alleles alone (i.e., the MLST phylogeny) did not mirror the WGS-based *B. cereus s.l.* phylogeny perfectly. Regardless of the ANI threshold used (i.e., 92.5, 94, 95, or 96 ANI), the *B. cereus s.l.* MLST phylogeny yielded polyphyletic genomospecies clusters (Figure 5 and Supplementary Figures S1-S5), although genomospecies clusters formed at 92.5 ANI reduced the proportion of polyphyletic genomospecies within the MLST phylogeny. One of eight genomospecies (12.5%) defined at 92.5 ANI were polyphyletic based on the MLST tree, compared to 2/11 (18.2%), 3/21 (14.3%), and 4/30 (13.3%) polyphyletic genomospecies defined at 94, 95, and 96 ANI respectively (Figure 5 and Supplementary Figures S1-S5).

### An adjusted, eight-group *panC* framework remains largely congruent with proposed *B. cereus s.l.* genomospecies definitions

Another popular typing method used to assign *B. cereus s.l.* isolates to major phylogenetic groups relies on the sequence of *panC* (Guinebretiere et al., 2008; Guinebretiere et al., 2010). However, the seven-group *panC* framework had to be adjusted to accommodate the growing amount of *B. cereus s.l.* genomic diversity provided by WGS, as *panC* sequences assigned to Groups II, III, and VI using the seven-group typing scheme implemented in the original BTyper were polyphyletic (Figure 4A).

The adjusted, eight-group *panC* framework constructed here (Figure 4B) and implemented in BTyper3 v. 3.1.0 resolved all polyphyletic *panC* group assignments (Figure 4). *panC* group assignments using the adjusted, eight-group framework described here, as well as those obtained using the original seven-group framework implemented in BTyper v. 2.3.3, are available for 2,229 *B. cereus s.l.* genomes (Table 1 and Supplementary Tables S1 and S6). Note that group assignments using the seven-group framework implemented in the web-tool published by Guinebretiere, et al. (Guinebretiere et al., 2010) are not available, as the database is not publicly available, and the web-based method is not scalable.

However, even with an improved eight-group framework for *panC* group assignment, the *B. cereus s.l. panC* phylogeny yielded polyphyletic genomospecies, regardless of the ANI-based threshold used to define genomospecies. For seven of the eight *B. cereus s.l.* genomospecies defined at 92.5 ANI (with the exclusion of effective and proposed putative species), the *panC* locus produced a monophyletic clade for each genomospecies (Figures 4 and 5 and Supplementary Figures S6 and S7). However, based on the sequence of *panC*, the *B. mosaicus* genomospecies was polyphyletic, with the *panC* sequence of *B. luti* forming a separate lineage within the *B. mosaicus panC* clade (Figure 5 and Supplementary Figures S6 and S7). Similarly, genomospecies defined at 94, 95, and 96 ANI produced even greater proportions of polyphyletic *panC* clusters, with 5/11 (45.5%), 8 or 9/21 (38.1 or 42.9%, depending on the phylogeny rooting method), and 8/30 (26.7%) genomospecies polyphyletic via *panC*, respectively (Supplementary Figures S6-S15).

### *rpoB* provides lower resolution than *panC* for single-locus sequence typing of *B. cereus s.l*. isolates

Another popular single-locus sequence typing method for characterizing spore-forming bacteria, including *B. cereus s.l.* isolates, relies on sequencing *rpoB*, which encodes the beta subunit of RNA polymerase (Huck et al., 2007b; Ivy et al., 2012). Among publicly available *B. cereus s.l.* isolate genomes, ATs assigned using the Cornell University Food Safety Laboratory and Milk Quality Improvement Program’s (CUFSL/MQIP) *rpoB* allelic typing database (Carroll et al., 2017), much like STs assigned using PubMLST’s seven-gene scheme (described above), were each confined to a single genomospecies at 92.5 ANI, with no AT split across genomospecies (Supplementary Tables S1 and S7). However, fewer than 2/3 of all *B. cereus s.l.* genomes possessed a *rpoB* allele that matched a member of the database exactly (i.e., with 100% nucleotide identity and coverage; 1,425/2,231 genomes, or 63.9%). Additionally, the *B. cereus s.l. rpoB* phylogeny showcased numerous polyphyletic genomospecies, regardless of the ANI threshold at which genomospecies were defined (3/8 [37.5%], 3 or 4/11 [27.2 or 36.4%, depending on the phylogeny rooting method], 6 or 9/21 [28.6 or 42.9%, depending on the phylogeny rooting method], and 8/30 [26.7%] polyphyletic *rpoB* clades among genomospecies defined at 92.5, 94, 95, and 96 ANI, respectively; Figure 5 and Supplementary Figures S16-S23).

### Numerous single loci mirror the topology of *B. cereus s.l.* and may provide improved resolution for single- and/or multi-locus sequence typing

A total of 1,719 single-copy loci were present among 313 high-quality *B. cereus s.l.* medoid genomes identified at 99 ANI (this was done to remove highly similar genomes and reduce the search space). After alignment, 255 of the 1,719 loci (i) produced an alignment that did not include any gap characters among at least 90% of its sites and (ii) contained a continuous sequence, uninterrupted by gaps, which covered at least 90% of total sites within the alignment, (iii) were present in a single copy in all 1,741 high-quality *B. cereus s.l.* genomes, sharing ≥ 90% nucleotide identity and coverage with at least one of the 313 alleles extracted from each of the 313 99 ANI medoid genomes, and (iv) produced a maximum likelihood phylogeny which mirrored the WGS phylogeny (Kendall-Colijn *P* < 0.05 after a Bonferroni correction; Supplementary Table S2). The resulting 255 single-copy core loci spanned a wide array of functions and were predicted to be involved in a diverse range of biological processes, including sporulation and response to stress (Figure 6, Supplementary Figure S24, and Supplementary Table S2).

**Figure 6.**
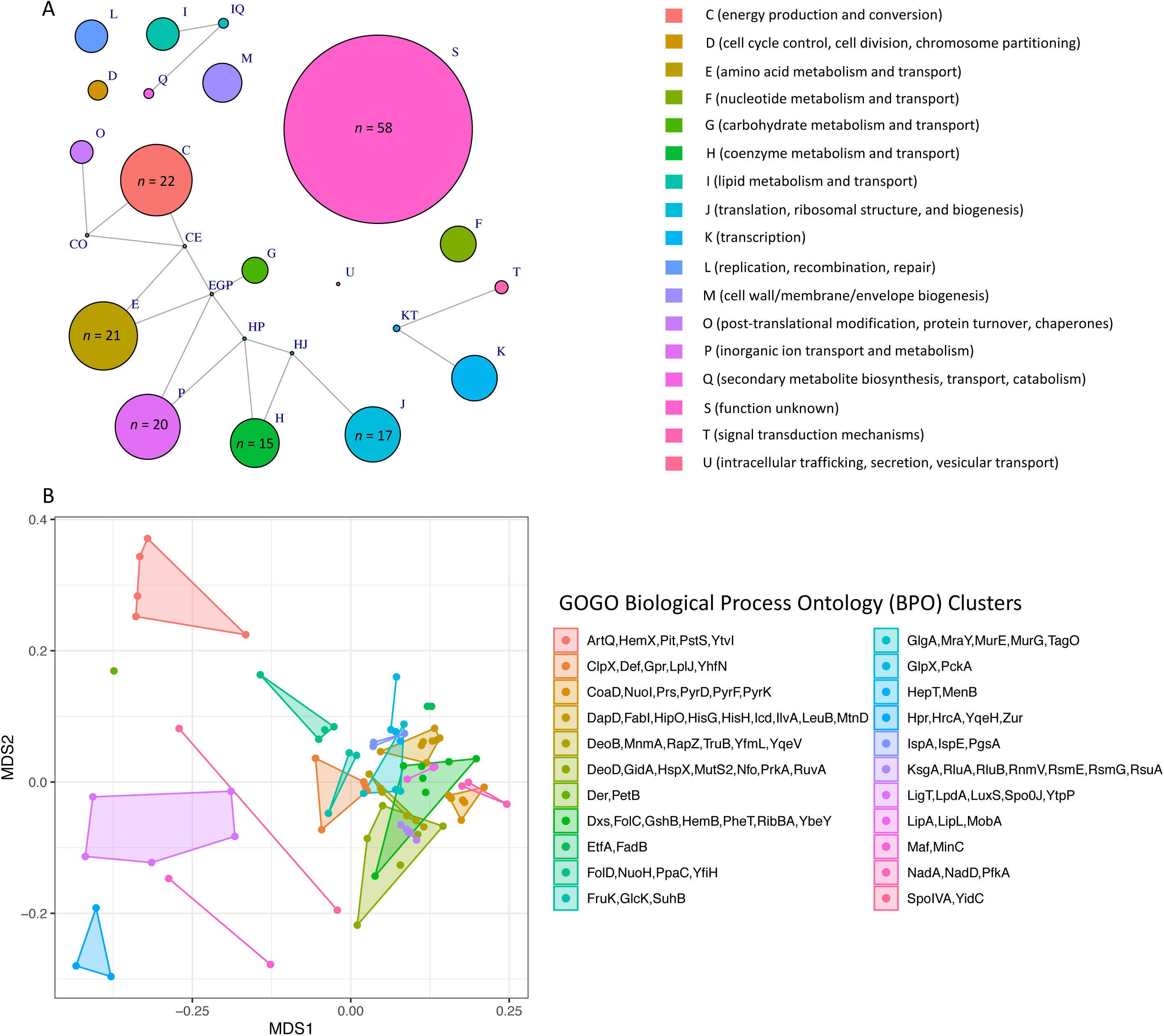
(A) Network of Clusters of Orthologous Groups (COG) functional categories assigned to 255 single-copy core genes that topologically mirror the *B. cereus s.l.* whole-genome phylogeny (Kendall-Colijn *P* < 0.05 after a Bonferroni correction). Each node corresponds to a COG functional category/group of functional categories assigned to one or more genes. Node size corresponds to the number of genes (out of 255 possible genes) assigned to a functional category/group of functional categories, ranging from one to 58 (for S, function unknown). Edges connect nodes that share one or more functional categories. Nodes of functional categories assigned to 15 or more genes are annotated with a text label denoting the number of genes assigned to the respective functional category. (B) Results of nonmetric multidimensional scaling (NMDS) performed using pairwise semantic/functional dissimilarities calculated between 94 single-copy core genes based on their assigned Gene Ontology (GO) Biological Process Ontology (BPO) terms. Points represent individual genes, while shaded regions and convex hulls correspond to clusters of genes identified by GOGO, based on their BPO similarities. For a complete list of annotations associated with each of the 255 single-copy core genes, see Supplementary Table S2. For NMDS plots constructed using Cellular Component Ontology (CCO) and Molecular Function Ontology (MFO) dissimilarities, see Supplementary Figure S24.

### The adjusted, eight-group *panC* framework captures genomic heterogeneity of anthrax-causing “*B. cereus*”

The set of 1,741 high-quality *B. cereus s.l.* genomes was queried for *B. cereus s.l.* virulence factors with known associations to anthrax (Okinaka et al., 1999; Candela and Fouet, 2006; Oh et al., 2011), emetic (Ehling-Schulz et al., 2006; Ehling-Schulz et al., 2015), and diarrheal illnesses (Schoeni and Wong, 2005; Stenfors Arnesen et al., 2008; Fagerlund et al., 2010; Senesi and Ghelardi, 2010) using amino acid identity and coverage thresholds of 70% and 80%, respectively (Figure 3). Using the proposed genomospecies-subspecies-biovar taxonomy and operon/cluster-level groupings for virulence factors (where applicable), cereulide synthetase-encoding *cesABCD* were detected in (i) the *B. mosaicus* and *B. mycoides* genomospecies and (ii) *panC* Group III and VI, respectively, as described previously (Guinebretiere et al., 2008; Guinebretiere et al., 2010; Carroll et al., 2017; Carroll and Wiedmann, 2020; Carroll et al., 2020) and regardless of whether the legacy seven-group or adjusted eight-group *panC* typing schemes were used (Figure 7 and Supplementary Table S1).

**Figure 7.**
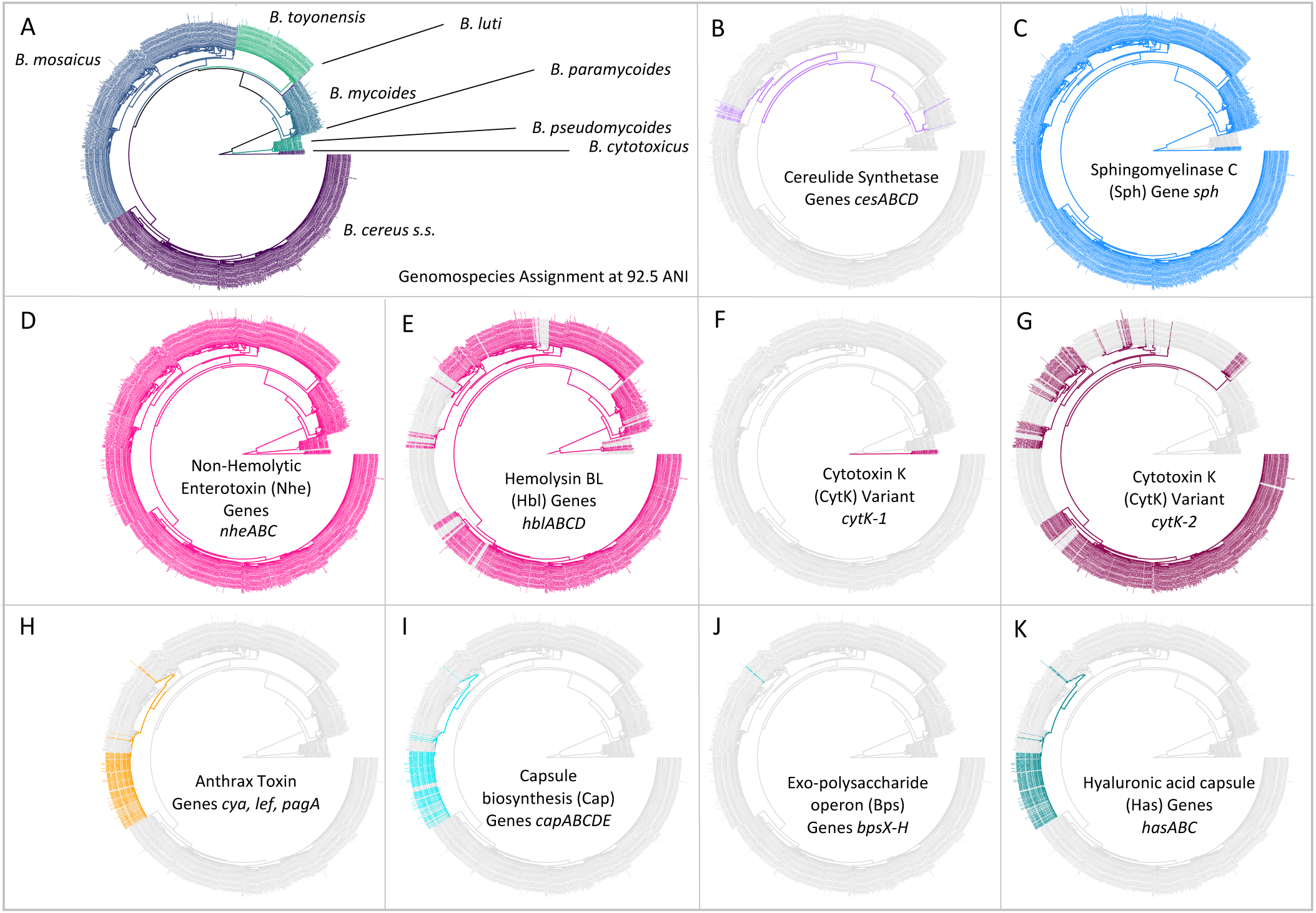
Distribution of selected *B. cereus s.l.* virulence factors within the *B. cereus s.l.* phylogeny (*n* = 1,741). Tip labels and branches within the phylogeny are colored by (A) *B. cereus s.l.* genomospecies, assigned using medoid genomes obtained at a 92.5 ANI threshold, and (B through K) presence and absence of the denoted *B. cereus s.l.* virulence factor (colored and gray tip labels, respectively). Virulence factors were detected using BTyper v. 3.1.0, with minimum amino acid identity and coverage thresholds of 70 and 80%, respectively, and a maximum E-value threshold of 1E-5. A virulence factor was considered to be present in a genome if all genes comprising the virulence factor were detected at the aforementioned thresholds; likewise, if one or more genes comprising a virulence factor were not detected at the given thresholds, the virulence factor was considered to be absent. The phylogeny was constructed using core SNPs identified in 79 single-copy orthologous gene clusters present among all 2,231 *B. cereus s.l.* genomes available in NCBI’s RefSeq database (accessed 19 November 2018; see Carroll, et al. 2020 for detailed methods) (Carroll et al., 2020). The type strain of “*B. manliponensis*” (i.e., the most distantly related member of the group) was treated as an outgroup on which the phylogeny was rooted. Tips representing genomes that (i) did not meet the quality thresholds and/or (ii) were not assigned to one of eight published genomospecies (i.e., genomes of unpublished, proposed, or effective *B. cereus s.l.* species) were omitted.

Anthrax toxin genes and anthrax-associated capsule-encoding operons *cap, has*, and *bps* were detected in their entirety in the *B. mosaicus* genomospecies alone (Figure 7 and Supplementary Table S1). Using the legacy, seven-group *panC* group assignment scheme implemented in the original BTyper (i.e., BTyper v. 2.3.3), all anthrax-associated virulence factors were confined to *panC* Group III; however, using the adjusted, eight-group framework, some anthrax-causing strains were assigned to Group II (Figure 7 and Supplementary Table S1). All anthrax-causing strains that belonged to the nonmotile, nonhemolytic (Tallent et al., 2012; Tallent et al., 2019) highly similar (≥99.9 ANI) (Jain et al., 2018; Carroll et al., 2020) lineage commonly associated with anthrax disease (known as species *B. anthracis*; using the proposed taxonomy, *B. anthracis* biovar Anthracis or *B. mosaicus* subsp. *anthracis* biovar Anthracis using subspecies and full notation, respectively) remained in *panC* Group III (Supplementary Table S1). However, the eight-group *panC* framework was able to capture genomic differences between anthrax-causing strains with phenotypic characteristics resembling *“B. cereus*” (e.g., motility, hemolysis; see Supplementary Table S1 here or Supplementary Table S5 of Carroll, et al. for a list of strains) (Carroll et al., 2020). Known previously as anthrax-causing “*B. cereus*” or “*B. cereus*” biovar Anthracis, among other names (using the proposed 2020 taxonomy, *B. mosaicus* biovar Anthracis), these strains could be partitioned into two major lineages: one that more closely resembled *B. anthracis* and one that more closely resembled *B. tropicus* using ANI-based comparisons to species type strains that existed in 2019 (Carroll et al., 2020). These anthrax-causing “*B. cereus*” lineages were assigned to *panC* Groups III and II using the adjusted, eight-group *panC* framework developed here, respectively (Supplementary Table S1).

Diarrheal enterotoxin-encoding genes were widespread throughout the *B. cereus s.l.* phylogeny (Figure 7 and Supplementary Table S1), as many others have noted before (Guinebretiere et al., 2008; Stenfors Arnesen et al., 2008; Guinebretiere et al., 2010; Kovac et al., 2016; Carroll et al., 2017). Nhe-encoding *nheABC* were detected in nearly all genomes (1,731/1,741 genomes, 99.4%; Figure 7 and Supplementary Table S1). Hbl-encoding *hblABCD* were detected in one or more members of all genomospecies except *B. cytotoxicus* and *B. luti* (Figure 7 and Supplementary Table S1). Variant 2 of CytK-encoding *cytK* (i.e., *cytK-2*) was identified in *B. cereus s.s.* (Group IV), *B. mosaicus* (Groups II and III), and *B. toyonensis* (Group V); variant 1 (*cytK-1*) was exclusive to *B. cytotoxicus*, as noted previously (Fagerlund et al., 2004; Guinebretiere et al., 2006; Carroll et al., 2017; Stevens et al., 2019).

### A method querying recent gene flow identifies multiple major gene flow units among the *B. cereus s.s., B. mosaicus, B. mycoides*, and *B. toyonensis* genomospecies

A recently proposed method for delineating microbial gene flow units using recent gene flow (referred to hereafter as the “populations as clusters of gene transfer”, or “PopCOGenT”, method) (Arevalo et al., 2019) was applied to the set of 313 high-quality *B. cereus s.l.* medoid genomes identified at 99 ANI. The PopCOGenT method identified a total of 33 “main clusters”, or gene flow units that attempt to mimic the classical species definition used for animals and plants (Table 2). Minimum ANI values shared between isolates assigned to the same gene flow unit ranged from 94.7-98.9 ANI for clusters containing more than one isolate (Table 2). A “pseudo-gene flow unit” assignment method was implemented in BTyper3 v. 3.1.0, in which ANI values are calculated between a query genome and the medoid genomes of the 33 PopCOGenT gene flow units using FastANI; if the query genome shares an ANI value with one of the gene flow unit medoid genomes that is greater than or equal to the previously observed ANI boundary for the gene flow unit, it is assigned to that particular pseudo-gene flow unit (Figures 1 and 2 and Table 2). This pseudo-gene flow unit assignment method was applied to all 2,231 *B. cereus s.l.* genomes (Table 2 and Supplementary Table S1), and was found to yield pseudo-gene flow units that were each encompassed within a single genomospecies and *panC* group (using the adjusted eight-group *panC* scheme developed here), with no pseudo-gene flow units split across multiple genomospecies/*panC* groups (Table 2). PopCOGenT identified multiple gene flow units among the *B. cereus s.s., B. mosaicus, B. mycoides*, and *B. toyonensis* genomospecies delineated at 92.5 ANI (*n* = 4, 16, 7, and 2 main clusters, respectively; Figure 8 and Supplementary Figures S25-S32).

**Table 2.**
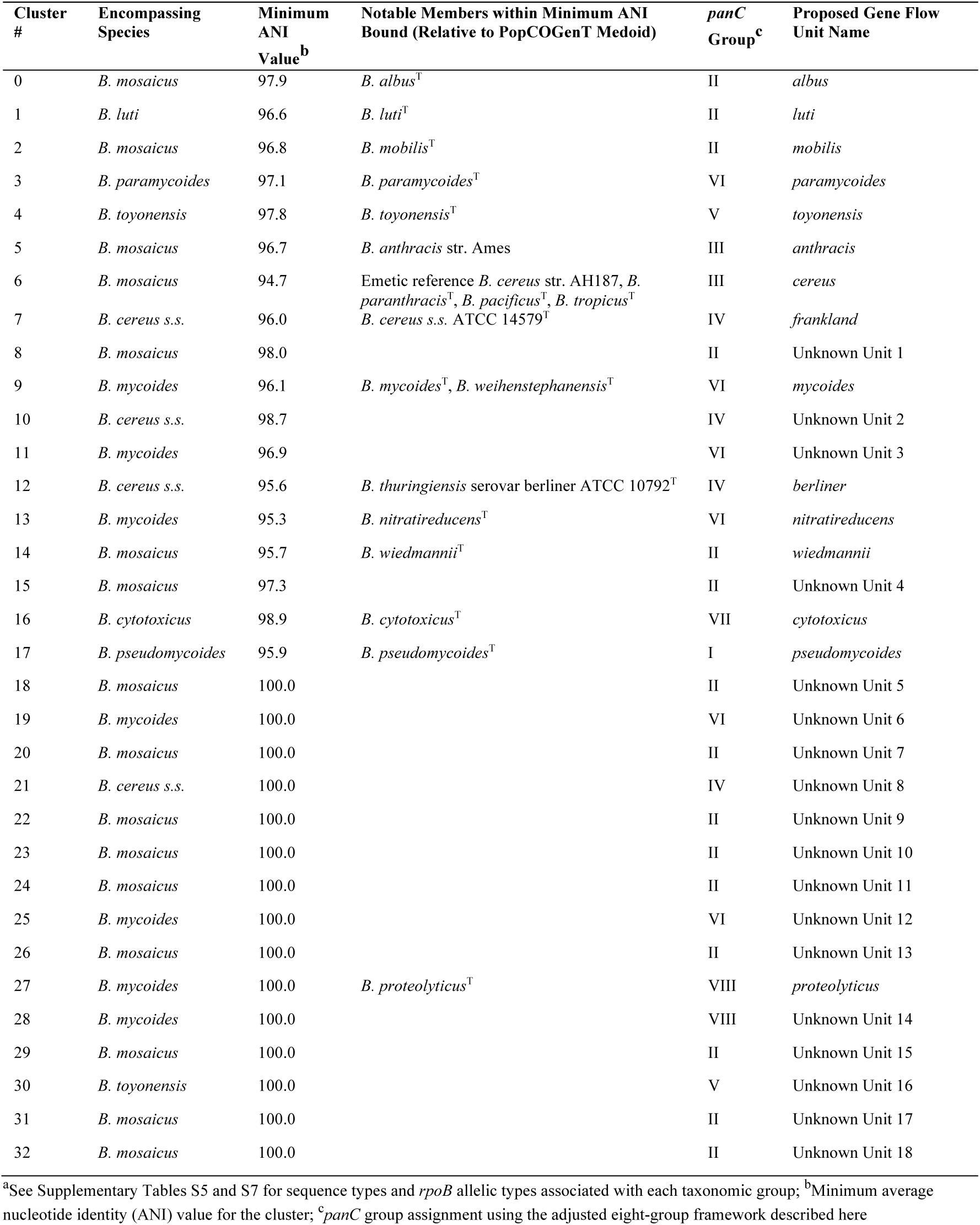
Gene flow units delineated using recent gene flow.^a^

**Figure 8.**
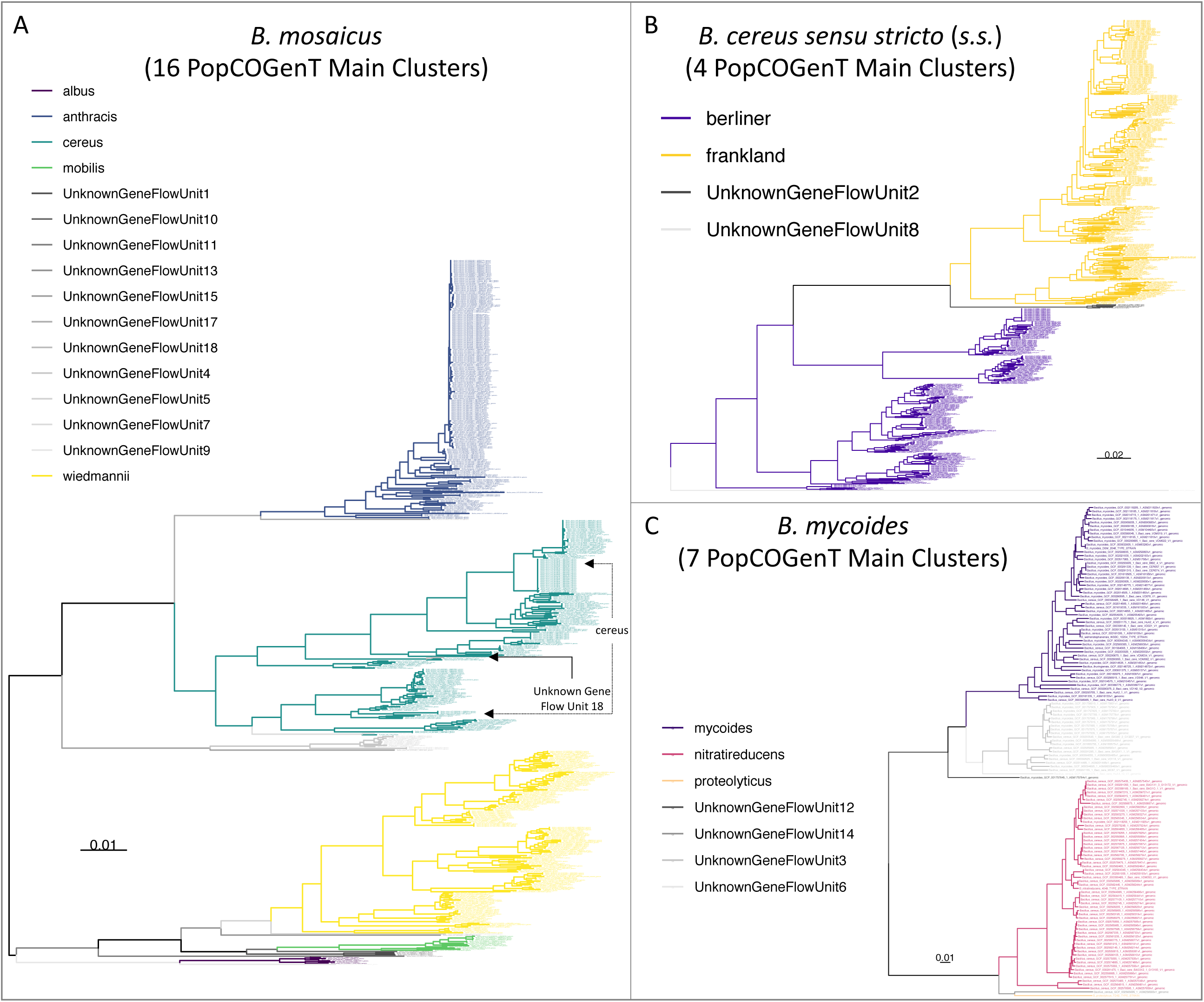
Maximum likelihood phylogenies constructed using genome-wide core SNPs identified among all high-quality genomes assigned to each of the (A) *B. mosaicus*, (B) *B. cereus sensu stricto* (*s.s*.), and (C) *B. mycoides* genomospecies delineated at a 92.5 ANI threshold. Branches and tip labels are colored by pseudo-gene flow unit assignment using the pseudo-gene flow unit assignment algorithm implemented in BTyper3 v. 3.1.0 (Figures 1 and 2). Only genomes that fell within the observed ANI boundary for each pseudo-gene flow unit are shown. Arrows are used to annotate polyphyletic pseudo-gene flow units derived from “true” gene flow units that also presented as polyphyletic (Supplementary Figure S25); the pseudo-gene flow units interspersed among them are additionally annotated with arrows. Phylogenies are rooted at the midpoint, and branch lengths are reported in substitutions per site. Genomospecies and pseudo-gene flow units were assigned using BTyper3 v. 3.1.0 and FastANI v. 1.0. For each phylogeny, core SNPs were identified among all high-quality genomes assigned to the genomospecies using kSNP3 v. 3.92 and the optimal *k*-mer size reported by Kchooser (*k* = 19 or 21). For each core SNP alignment, IQ-TREE v. 1.5.4 was used to construct a phylogeny, using the GTR+G+ASC nucleotide substitution model.

## Discussion

### The proposed *B. cereus s.l.* taxonomy is backwards-compatible with *B. cereus s.l.* species defined using historical ANI-based species thresholds

ANI-based methods have been used to define 12 *B. cereus s.l.* species prior to 2020: *B. cytotoxicus* and *B. toyonensis*, each proposed as novel species in 2013 (Guinebretiere et al., 2013; Jimenez et al., 2013), *B. wiedmannii* (proposed as a novel species in 2016) (Miller et al., 2016), and nine species (*B. albus, B. luti, B. mobilis, B. nitratireducens, B. pacificus, B. paranthracis, B. paramycoides, B. proteolyticus*, and *B. tropicus*) proposed in 2017 (Liu et al., 2017). However, the lack of a standardized ANI-based genomospecies threshold for defining *B. cereus s.l.* genomospecies has led to confusion regarding how *B. cereus s.l.* species should be delineated. *B. toyonensis* and the nine species proposed in 2017, for example, were defined using genomospecies thresholds of 94 and 96 ANI, respectively (Jimenez et al., 2013; Liu et al., 2017). The descriptions of *B. cytotoxicus* and *B. wiedmannii* as novel species each explicitly state that a 95 ANI threshold was used (Guinebretiere et al., 2013; Miller et al., 2016); however, the *B. wiedmannii* type strain genome shared a much higher degree of similarity with the type strain genome of its neighboring species than did *B. cytotoxicus* (Miller et al., 2016). As such, choice of ANI-based genomospecies threshold can affect which *B. cereus s.l.* strains belong to which genomospecies, and may even produce overlapping genomospecies in which a genome can belong to more than one genomospecies (Carroll et al., 2020).

The proposed *B. cereus s.l.* taxonomy (Carroll et al., 2020) provides a standardized genomospecies threshold of 92.5 ANI, which has been shown to yield non-overlapping, monophyletic *B. cereus s.l.* genomospecies. However, the practice of assigning *B. cereus s.l.* isolates to genomospecies using species type strain genomes and historical species thresholds (i.e., 94-96 ANI) has been important for whole-genome characterization for *B. cereus s.l.* strains, including those responsible for illnesses and/or outbreaks (Lazarte et al., 2018; Bukharin et al., 2019; Carroll et al., 2019). Here, we show that all 18 published *B. cereus s.l.* genomospecies defined prior to 2020 are safely integrated into the proposed *B. cereus s.l.* taxonomy without polyphyly, regardless of whether a 94, 95, or 96 ANI genomospecies threshold was used to delineate species relative to type strain genomes.

### Single- and multi-locus sequence typing methods can be used to assign *B. cereus s.l.* isolates to species within the proposed taxonomy

Single- and multi-locus sequence typing approaches have been—and continue to be—important methods for classifying *B. cereus s.l.* isolates into phylogenetic units. They have been used to characterize *B. cereus s.l.* strains associated with illnesses and outbreaks (Cardazzo et al., 2008; Glasset et al., 2016; Akamatsu et al., 2019; Carroll et al., 2019), strains isolated from food and food processing environments (Huck et al., 2007a; Thorsen et al., 2015; Kindle et al., 2019; Ozdemir and Arslan, 2019; Zhuang et al., 2019; Zhao et al., 2020), and strains with industrial applications (e.g., biopesticide strains) (Johler et al., 2018). Additionally, STs and ATs assigned using these approaches have been used to construct frameworks for predicting the risk that a particular *B. cereus s.l.* strain poses to food safety, public health, and food spoilage (Guinebretiere et al., 2010; Rigaux et al., 2013; Buehler et al., 2018; Miller et al., 2018; Webb et al., 2019). It is thus important to ensure that the proposed standardized taxonomy for *B. cereus s.l.* remains congruent with widely used sequence typing approaches.

Here, we assessed the congruency of three popular single- and multi-locus sequence typing schemes for *B. cereus s.l.* with proposed genomospecies definitions: (i) the PubMLST seven-gene MLST scheme for *B. cereus s.l.* (Jolley and Maiden, 2010; Jolley et al., 2018), (ii) the seven-group *panC* typing scheme developed by Guinebretierere, et al. (Guinebretiere et al., 2008; Guinebretiere et al., 2010) as implemented in the original BTyper (Carroll et al., 2017), and (iii) the CUFSL/MQIP *rpoB* allelic typing scheme used for characterizing spore-forming bacteria, including members of *B. cereus s.l.* (Durak et al., 2006; Huck et al., 2007a; Ivy et al., 2012; Buehler et al., 2018). STs and ATs assigned using MLST and *rpoB* allelic typing, respectively, as well as six of eight *panC* groups assigned using the adjusted eight-group framework developed here, were each contained within a single genomospecies at 92.5 ANI. Thus, past studies employing these methods can be easily interpreted within the proposed taxonomic framework for the group.

MLST, *panC* group assignment, and *rpoB* allelic typing will likely remain extremely valuable for characterizing *B. cereus s.l.* isolates, as all three approaches remain largely congruent with *B. cereus s.l.* genomospecies defined at 92.5 ANI. However, all three typing methods produced at least one polyphyletic genomospecies among genomospecies defined at 92.5 ANI. Higher, historical genomospecies thresholds (i.e., 94, 95, and 96 ANI) showcased even higher proportions of polyphyly within the MLST and *panC* phylogenies. This observation is particularly important for *panC* group assignment, as *panC* may not be able to differentiate between some members of *B. mosaicus* and *B. luti* (each assigned to *panC* Group II) with adequate resolution. In addition to assessing the congruency of proposed typing methods, we used a computational approach to identify putative loci that may better capture the topology of the whole-genome *B. cereus s.l.* phylogeny. While typing schemes that incorporate these loci still need to be validated in an experimental setting, future single-locus sequence typing methods using loci that mirror the “true” topology of *B. cereus s.l.* may improve sequence typing efforts.

### A rapid, scalable ANI-based method can be used to assign genomes to pseudo-gene flow units identified among *B. cereus s.l.* genomospecies

ANI-based methods have become the gold standard for bacterial taxonomy in the WGS era (Richter and Rossello-Mora, 2009), as they conceptually mirror DNA-DNA hybridization and implicitly account for the fluidity that accompanies bacterial genomes (Jain et al., 2018). However, the concept of the bacterial “species” has been, and remains, controversial, as the promiscuous genetic exchange that occurs among prokaryotes can obscure population boundaries (Hanage et al., 2005; Rocha, 2018; Arevalo et al., 2019). Recently, Arevalo, et al. (Arevalo et al., 2019) outlined a method that attempts to delineate microbial gene flow units and the populations within them using a metric based on recent gene flow. The resulting gene flow units identified among bacterial genomes are proposed to mimic the classical species definition used for plants and animals (i.e., interbreeding units separated by reproductive barriers) (Huxley, 1943; Arevalo et al., 2019). Here, we used PopCOGenT to characterize a subset of isolates that capture genomic diversity across *B. cereus s.l.*, and we identified 33 main gene flow units among *B. cereus s.l.* isolates assigned to known genomospecies.

While the PopCOGenT method attempts to apply classical definitions of species developed with higher organisms in mind to microbes, we propose to maintain ANI-based *B. cereus s.l.* genomospecies definitions (i.e., ANI-based genomospecies clusters formed using medoid genomes obtained at a 92.5 ANI breakpoint) due to (i) the speed, scalability, portability, and accessibility of the ANI algorithm, and (ii) the accessibility and backwards-compatibility of the eight-genomospecies *B. cereus s.l.* taxonomic framework, as demonstrated in this study. ANI is fast and can readily scale to large numbers (e.g., tens of thousands) of bacterial genomes (Jain et al., 2018), traits that will become increasingly important as more *B. cereus s.l.* genomes are sequenced. In addition to speed and scalability, ANI is a well-understood algorithm implemented in many easily accessible tools, including command-line tools (e.g., FastANI, pyani, OrthoANI), desktop applications (e.g., JSpecies, OrthoANI), and web-based tools (e.g., JSpeciesWS, MiGA, OrthoANIu) (Goris et al., 2007; Richter and Rossello-Mora, 2009; Lee et al., 2016; Pritchard et al., 2016; Richter et al., 2016; Yoon et al., 2017; Jain et al., 2018; Rodriguez et al., 2018). Finally, the gene flow units identified using the PopCOgenT method in the present study were not congruent with historical ANI-based genomospecies assignment methods used for *B. cereus s.l.* Genomospecies defined at historical ANI thresholds are not readily integrated into the gene flow units identified via the PopCOgenT method, as the ANI boundaries for PopCOgenT gene flow units vary (Table 2).

Despite its infancy and current limitations, the PopCOgenT framework provides an interesting departure from a one-threshold-fits-all ANI-based taxonomy. Here, we implemented a “pseudo-gene flow unit” method in BTyper3 v. 3.1.0 that can be used to assign a user’s genome of interest to a pseudo-gene flow unit using the set of 33 PopCOgenT gene flow unit medoid genomes, the pairwise ANI values calculated within PopCOgenT gene flow units, and FastANI. However, it is essential to note the limitations of the pseudo-gene flow unit assignment method implemented in BTyper3. First and foremost, ANI and the methods employed by PopCOgenT are fundamentally and conceptually different; the pseudo-gene flow unit assignment method described here does not infer recent gene flow, nor does it use PopCOgenT or any of its metrics. Thus, the pseudo-gene flow unit assignment method cannot be used to construct true gene flow units for *B. cereus s.l*. Secondly, to increase the speed of PopCOGenT, we reduced *B. cereus s.l.* to a set of 313 representative genomes that encompassed the diversity of the species complex; genomes that shared ≥ 99 ANI with one or more genomes in this representative set were omitted (i.e., 1,428 of 1,741 high-quality genomes were omitted; 82.0%). Consequently, gene flow among most closely related lineages that shared ≥ 99 ANI with each other was not assessed, as it was thereby assumed that highly similar genomes that shared ≥ 99 ANI with each other belonged to the same PopCOGenT “main cluster” (i.e., “species”). It is possible that the inclusion of these highly similar genomes would have resulted in the discovery of additional gene flow units, or perhaps changes in existing ones, and future studies are needed to assess and refine this. However, the pseudo-gene flow unit assignment approach described here allows researchers to rapidly identify the most similar medoid genome of true gene flow units identified within *B. cereus s.l.* Results should not be interpreted as an assessment of recent gene flow, but rather as a higher-resolution phylogenomic clade assignment, similar to how one might use MLST for delineation of lineages within species. We anticipate that our rapid method will be valuable to researchers who desire greater resolution than what is provided at the genomospecies level, particularly when querying diverse *B. cereus s.l.* genomospecies that comprise multiple major clades (e.g., *B. mosaicus, B. mycoides, B. cereus s.s.*).

## Author Contributions

LMC performed all computational analyses. LMC, RAC, and JK designed the study and co-wrote the manuscript.

