## Supplementary Figure S9 for "No Assembly Required: Using BTyper3 to Assess the Congruency of a Proposed Taxonomic Framework for the *Bacillus cereus* group with Historical Typing Methods"

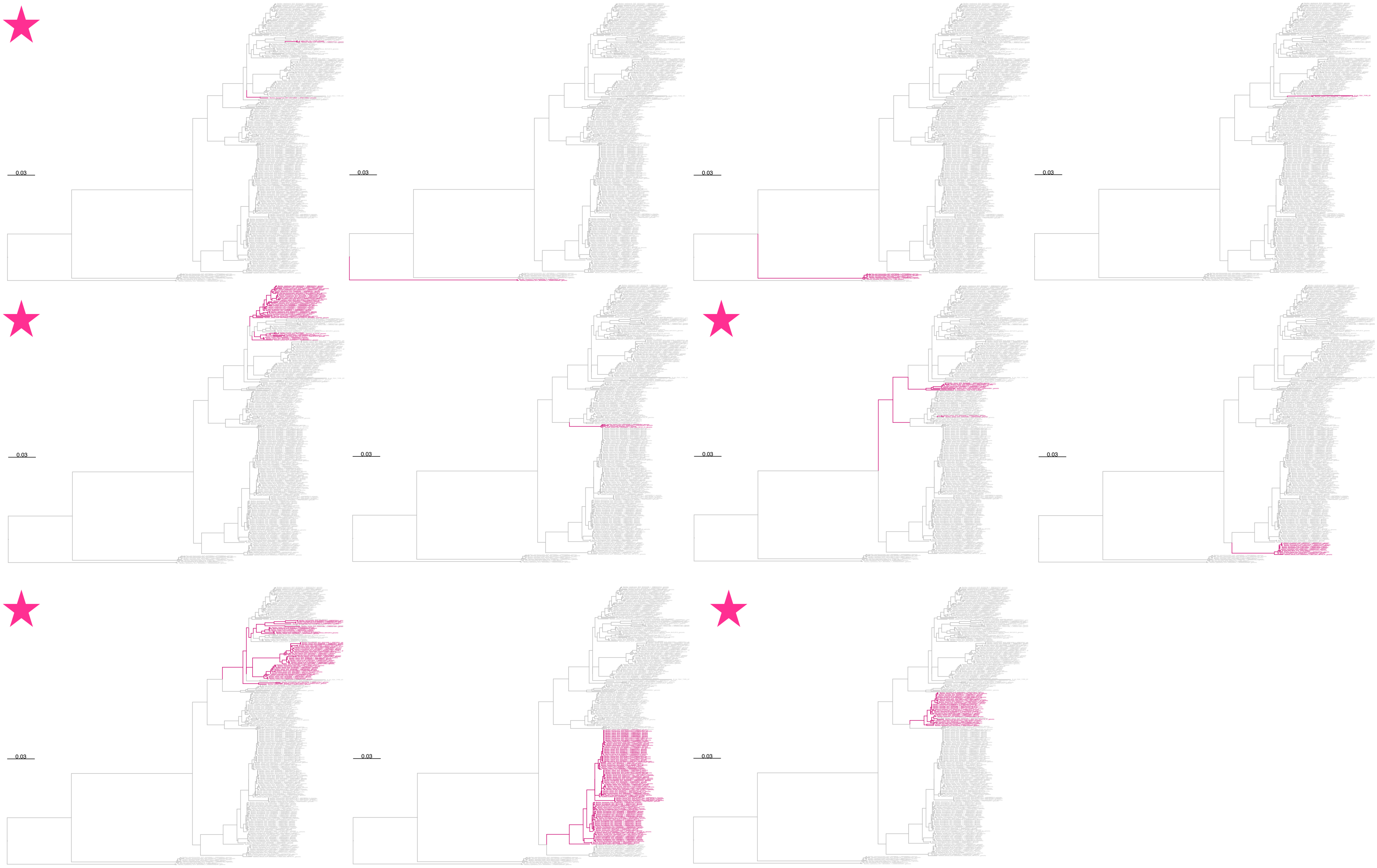


**Supplementary Figure S9.** Maximum likelihood phylogeny constructed using *panC*, extracted from 314 high-quality *B. cereus s.l.* genomes. Branches and tip labels are colored by ANI-based genomospecies assignment using medoid genomes identified among a set of 1,742 high-quality *B. cereus s.l.* genomes at 94 ANI; for each of the 11 genomospecies clusters, the "foreground" genomospecies is colored (pink) and the 10 background genomospecies are shown in gray. Phylogenies for which the foreground genomospecies (pink) presents as polyphyletic are annotated with a pink star in the upper left corner of the panel. Phylogenies are rooted using the *panC* sequence of the "*B. manliponensis*" type strain genome as an outgroup (not shown for readability), and branch lengths are reported in substitutions per site. High-quality genomes were derived from the total set of 2,231 *B. cereus s.l.* genomes available in NCBI's RefSeq database (accessed November 19, 2018) and met all of the following conditions ($n$ = 1,742)*:* (i) possessed an N50 > 100 kbp (via QUAST v. 4.0); (ii) possessed a CheckM "Completeness" score ≥ 97.5 (via CheckM v. 1.0.7); (iii) possessed a CheckM "Contamination" score ≤ 2.5; and (iv) belonged to one of eight published *B. cereus s.l.* genomospecies or "*B. manliponensis*" (i.e., effective/proposed *B. cereus s.l.* species "*B. bingmayongensis*", "*B. clarus*", and "*B. gaemokensis*", and putative novel species "Unknown Species 13-18" described by Carroll, et al. 2020 were not included). Medoid genomes were identified at a 94 ANI threshold using the bactaxR package in R version 3.6.1, and FastANI v. 1.0 was used to assign each of the 1,742 *B. cereus s.l.* genomes to a genomospecies cluster. To remove highly similar genomes for readability, medoid genomes ($n$ = 314) were identified among the set of 1,742 high-quality genomes at a 99 ANI clustering threshold using bactaxR. BTyper v. 2.3.3 was used to extract *panC* from each of the 314 *B. cereus s.l.* genomes, and MAFFT v. 7.453-with-extensions was used to construct an alignment. IQ-TREE v. 1.6.5 was used to construct a phylogeny, using the optimal nucleotide substitution model selected using ModelFinder (i.e., the TVM+F+R4 model).
