## Supplementary Figure S17 for "No Assembly Required: Using BTyper3 to Assess the Congruency of a Proposed Taxonomic Framework for the *Bacillus cereus* group with Historical Typing Methods"

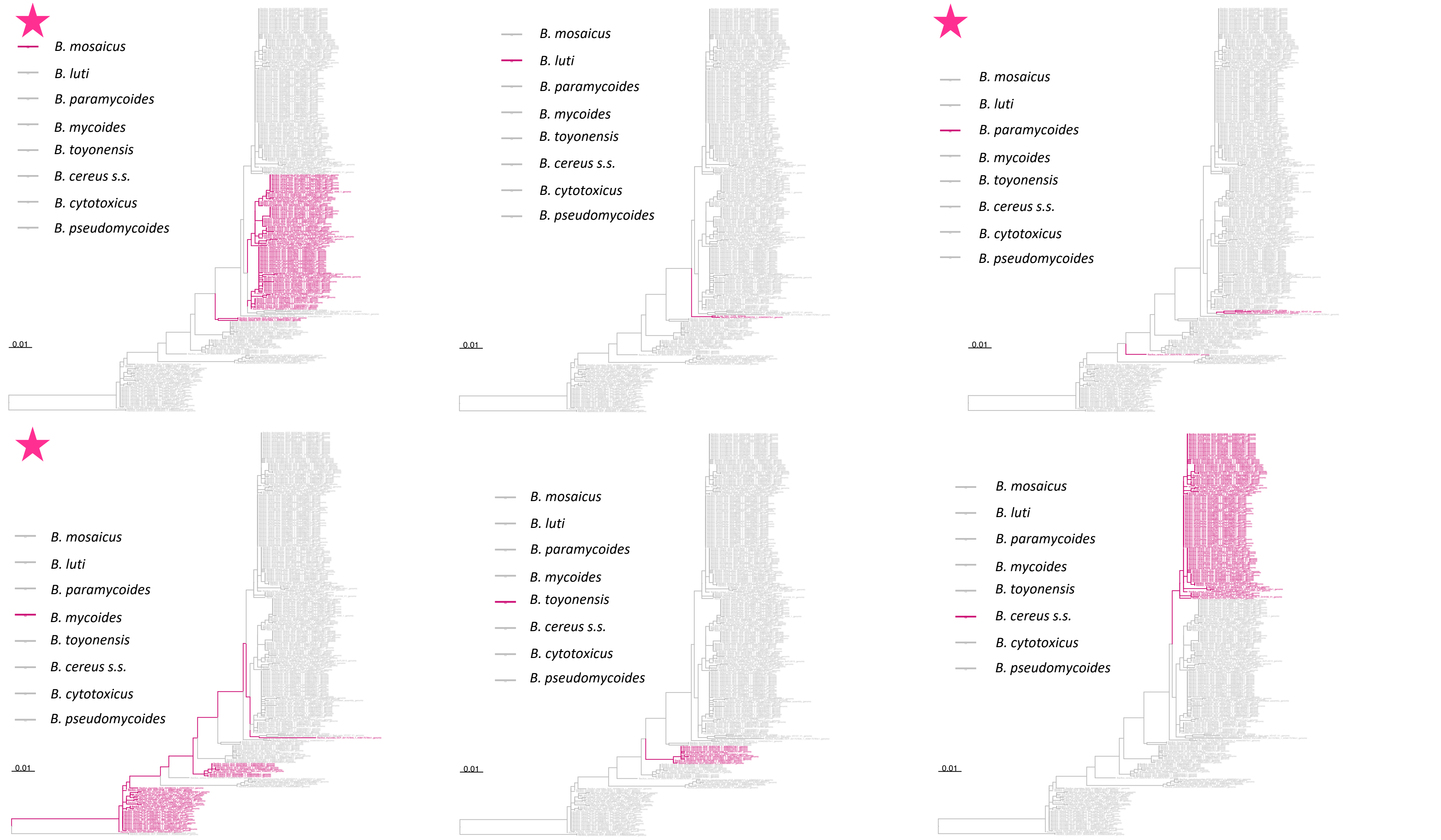

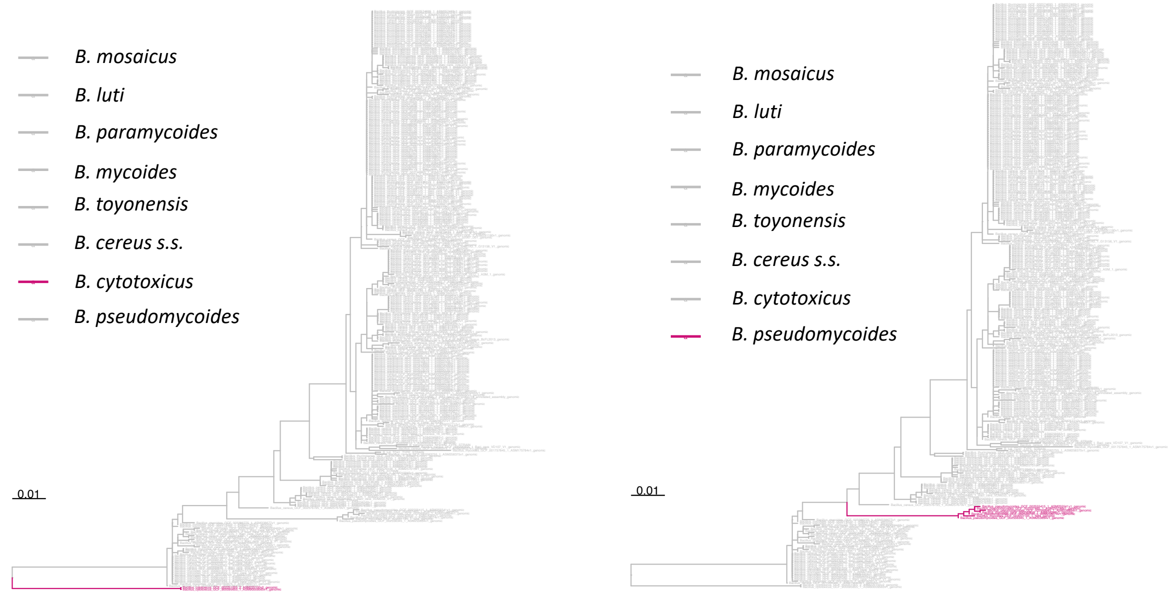


**Supplementary Figure S17.** Maximum likelihood phylogeny constructed using *rpoB*, extracted from 314 high-quality *B. cereus s.l.* genomes. Branches and tip labels are colored by ANI-based genomospecies assignment using the proposed *B. cereus s.l.* taxonomic framework (i.e., eight genomospecies assigned using medoid genomes and a 92.5 ANI threshold); for each of the eight genomospecies clusters, the “foreground” species is colored (pink) and the seven background species are shown in gray. Phylogenies for which the foreground genomospecies (pink) presents as polyphyletic are annotated with a pink star in the upper left corner of the panel. Phylogenies are rooted using the *rpoB* sequence of the “*B. manliponensis*” type strain genome as an outgroup (not shown for readability), and branch lengths are reported in substitutions per site. High-quality genomes were derived from the total set of 2,231 *B. cereus s.l.* genomes available in NCBI’s RefSeq database (accessed November 19, 2018) and met all of the following conditions ($n$ = 1,742): (i) possessed an N50 > 100 kbp (via QUAST v. 4.0); (ii) possessed a CheckM “Completeness” score ≥ 97.5 (via CheckM v. 1.0.7); (iii) possessed a CheckM “Contamination” score ≤ 2.5; and (iv) belonged to one of eight published *B. cereus s.l.* genomospecies or “*B. manliponensis*” (i.e., effective/proposed *B. cereus s.l.* species “*B. bingmayongensis*”, “*B. clarus*”, and “*B. gaemokensis*”, and putative novel species “Unknown Species 13-18” described by Carroll, et al. 2020 were not included). To remove highly similar genomes for readability, medoid genomes ($n$ = 314) were identified among the set of 1,742 high-quality genomes at a 99 ANI clustering threshold using the bactaxR package in R version 3.6.1. Genomospecies were assigned using BTyper3 v. 3.0.2 and FastANI v. 1.0. BTyper v. 2.3.3 was used to extract *rpoB* from each of the 314 *B. cereus s.l.* genomes, and MAFFT v. 7.453-with-extensions was used to construct an alignment. IQ-TREE v. 1.6.5 was used to construct a phylogeny, using the optimal nucleotide substitution model selected using ModelFinder (i.e., the TIM3+F+I+G4 model).
