## Supplementary Figure S24 for "No Assembly Required: Using BTyper3 to Assess the Congruency of a Proposed Taxonomic Framework for the *Bacillus cereus* group with Historical Typing Methods"

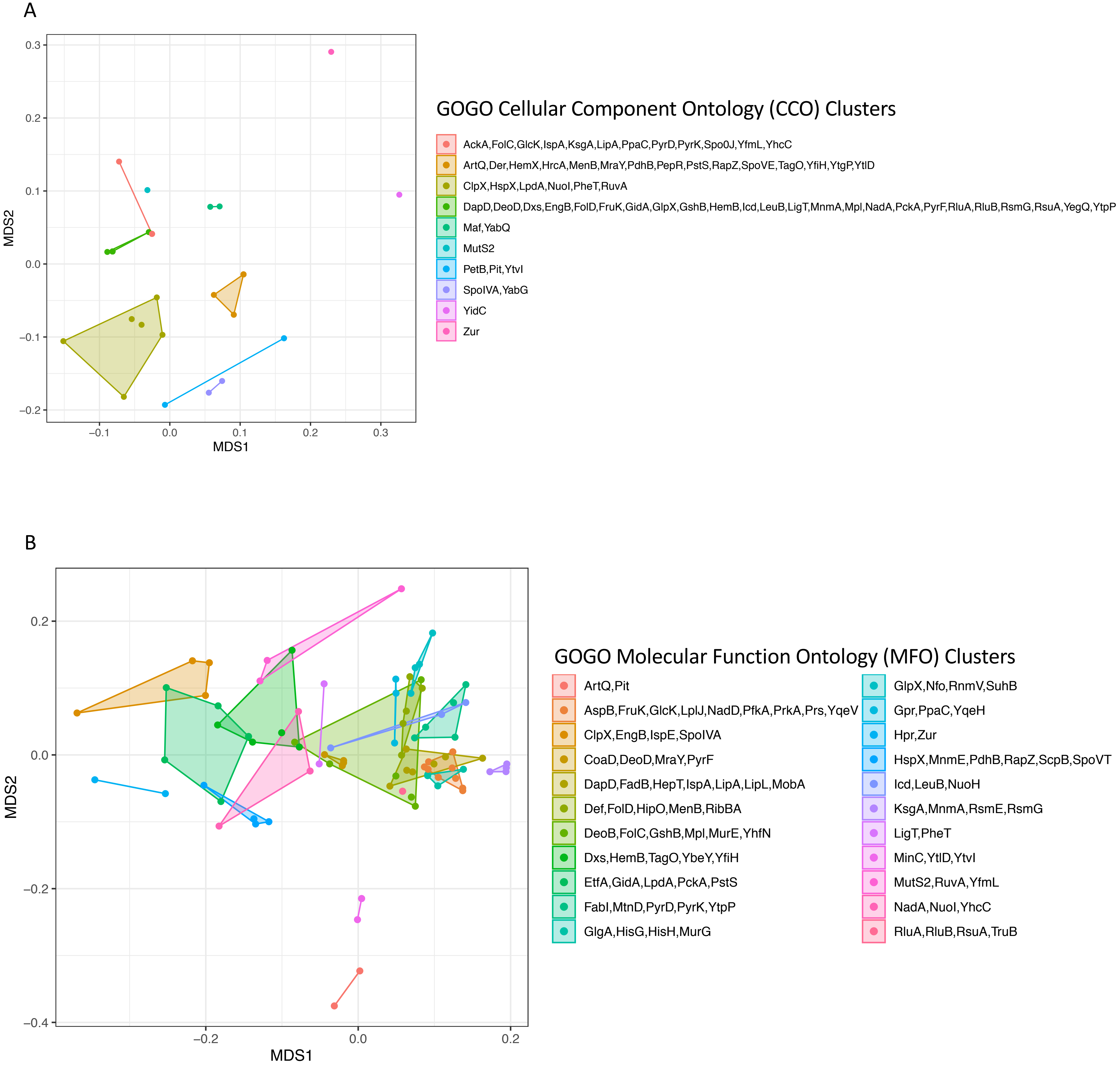


**Supplementary Figure S24.** Results of nonmetric multidimensional scaling (NMDS) performed using pairwise semantic/functional dissimilarities calculated between (A) 67 and (B) 93 single-copy core genes based on their assigned Gene Ontology (GO) (A) Cellular Component Ontology (CCO) and (B) Molecular Function Ontology (MFO) terms. Points represent individual genes, while shaded regions and convex hulls correspond to clusters of genes identified by GOGO, based on their (A) CCO and (B) MFO similarities. For a complete list of annotations associated with each of the 255 single-copy core genes, see Supplementary Table S2.
