## Supplementary Figure S25 for "No Assembly Required: Using BTyper3 to Assess the Congruency of a Proposed Taxonomic Framework for the *Bacillus cereus* group with Historical Typing Methods"

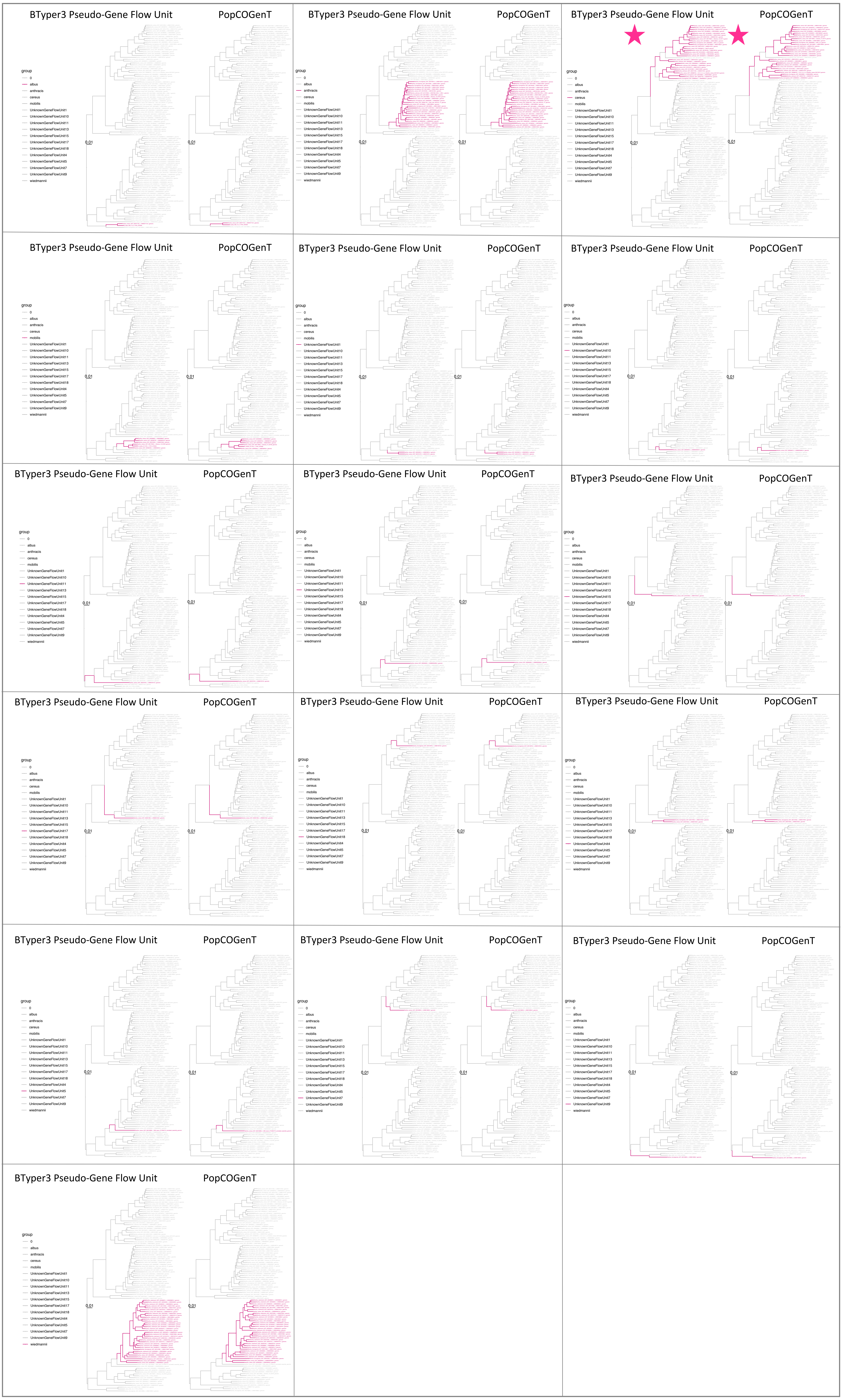


**Supplementary Figure S25.** Maximum likelihood phylogeny constructed using genome-wide core SNPs identified among all high-quality genomes assigned to the *B. mosaicus* genomospecies delineated at a 92.5 average nucleotide identity (ANI) threshold. Only tips of genomes that were identified as medoid genomes at a 99 ANI threshold are shown; this was done to reduce the set of 1,741 high-quality genomes to a set of 313 genomes that spanned *B. cereus s.l.* in its entirety which could then be queried using PopCOGenT in a computationally tractable amount of time. Each "main cluster" (i.e., "true" gene flow unit) identified by PopCOGenT is shown alongside its pseudo-gene flow unit identified using the pseudo-gene flow unit assignment method implemented in BTyper3 v. 3.1.0 (pink branches and tip labels), with background pseudo-gene flow units and true gene flow units denoted using gray tip labels and branches. Phylogenies in which the pseudo-gene flow unit and/or true gene flow unit present as polyphyletic are annotated with a pink star. Phylogenies are rooted at the midpoint, and branch lengths are reported in substitutions per site. Genomospecies and pseudo-gene flow units were assigned using BTyper3 v. 3.1.0 and FastANI v. 1.0. Core SNPs were identified among all high-quality *B. mosaicus* genomes using kSNP3 v. 3.92 and the optimal *k*-mer size reported by Kchooser (*k* = 19). IQ-TREE v. 1.5.4 was used to construct the phylogeny, using the resulting core SNP alignment and the GTR+G+ASC nucleotide substitution model.
