## Supplementary Figure S26 for "No Assembly Required: Using BTyper3 to Assess the Congruency of a Proposed Taxonomic Framework for the *Bacillus cereus* group with Historical Typing Methods"

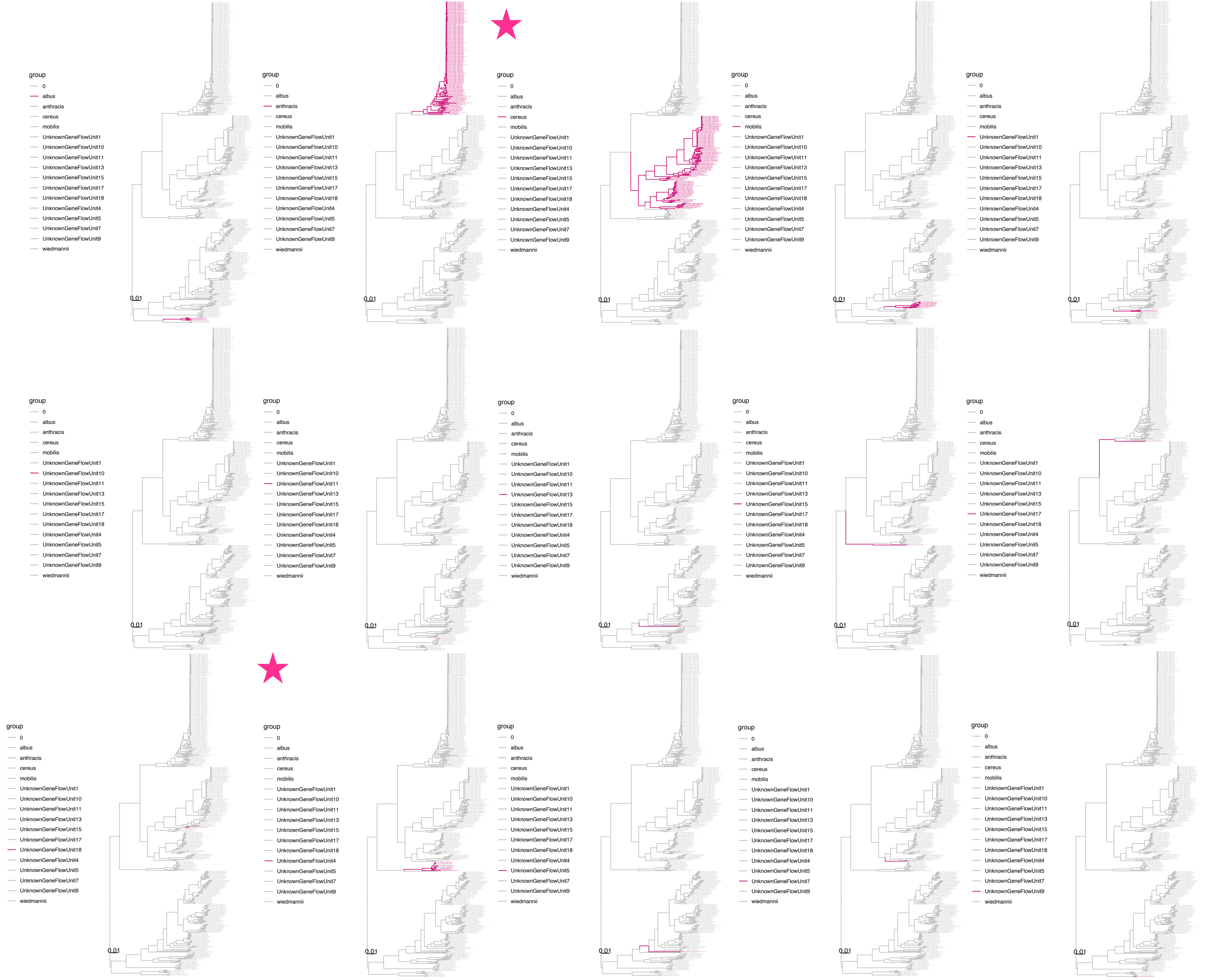


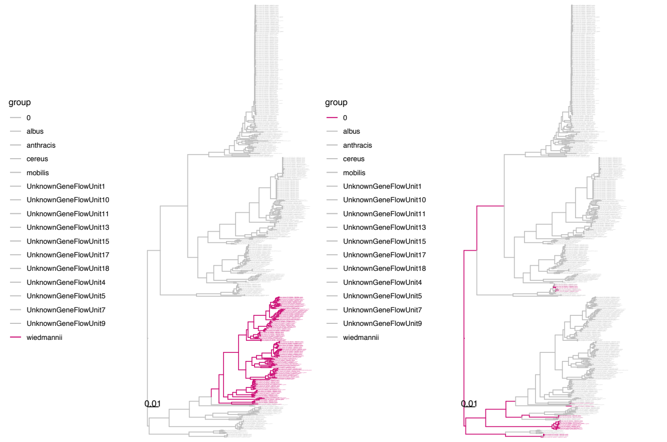


**Supplementary Figure S26.** Maximum likelihood phylogeny constructed using genome-wide core SNPs identified among all high-quality genomes assigned to the *B. mosaicus* genomospecies delineated at a 92.5 average nucleotide identity (ANI) threshold. Each pseudo-gene flow unit identified using the pseudo-gene flow unit assignment method implemented in BTyper3 v. 3.1.0 is shown in the foreground (pink branches and tip labels), with background pseudo-gene flow units denoted using gray tip labels and branches. Group 0 corresponds to genomes that did not fall within the observed ANI boundary for any gene flow unit. Phylogenies in which the pseudo-gene flow unit present as polyphyletic (excluding genomes that were not within the observed ANI boundaries of any gene flow unit) are annotated with a pink star. Phylogenies are rooted at the midpoint, and branch lengths are reported in substitutions per site. Genomospecies and pseudo-gene flow units were assigned using BTyper3 v. 3.1.0 and FastANI v. 1.0. Core SNPs were identified among all high-quality *B. mosaicus* genomes using kSNP3 v. 3.92 and the optimal *k*-mer size reported by Kchooser ($k$ = 19). IQ-TREE v. 1.5.4 was used to construct the phylogeny, using the resulting core SNP alignment and the GTR+G+ASC nucleotide substitution model.
