## Supplementary Figure S31 for "No Assembly Required: Using BTyper3 to Assess the Congruency of a Proposed Taxonomic Framework for the *Bacillus cereus* group with Historical Typing Methods"

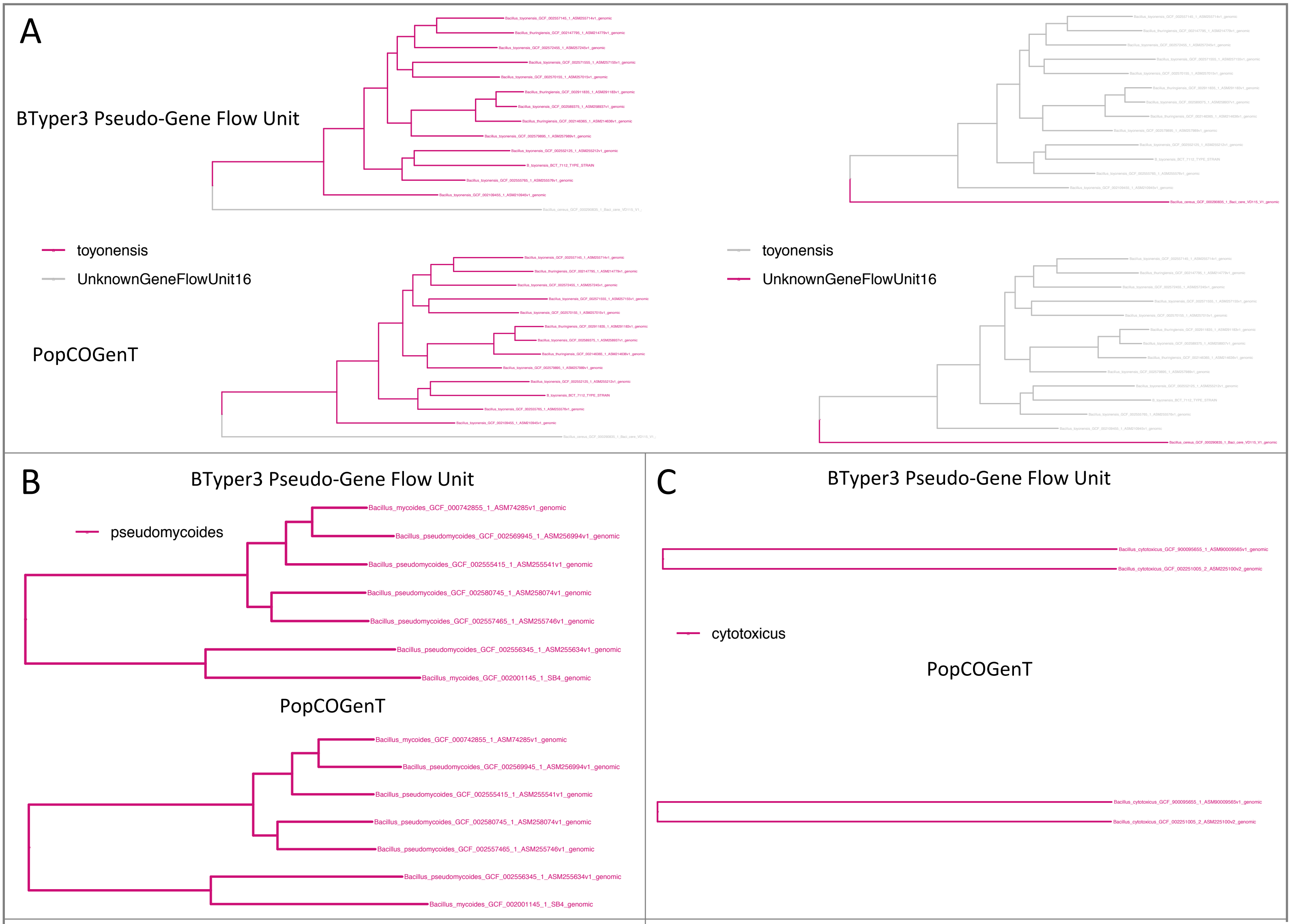

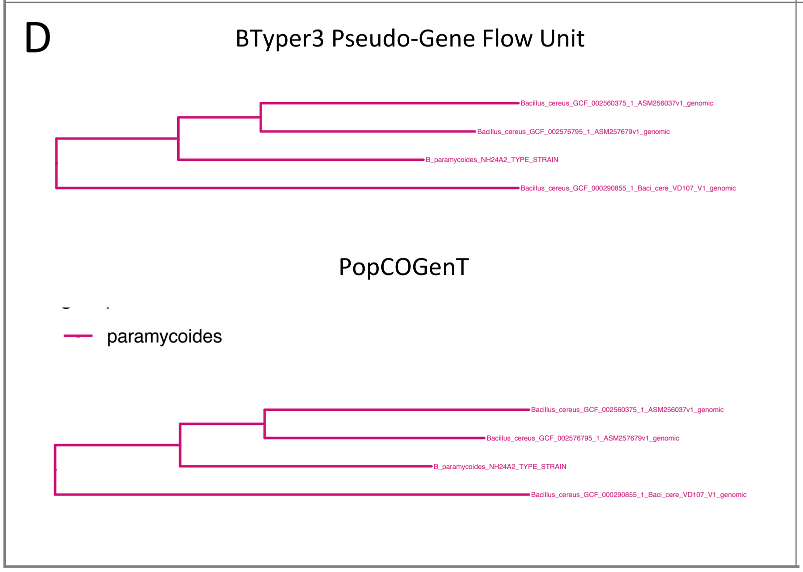


**Supplementary Figure S31.** Maximum likelihood phylogenies constructed using genome-wide core SNPs identified among all high-quality genomes assigned to the (A) *B. toyonensis*, (B) *B. pseudomycoides*, (C) *B. cytotoxicus*, and (D) *B. paramycoides* genomospecies delineated at a 92.5 average nucleotide identity (ANI) threshold. *B. luti* is not shown, as too few genomes were available to construct a phylogeny. Only tips of genomes that were identified as medoid genomes at a 99 ANI threshold are shown; this was done to reduce the set of 1,741 high-quality genomes to a set of 313 genomes that spanned *B. cereus s.l.* in its entirety which could then be queried using PopCOGenT in a computationally tractable amount of time. Each "main cluster" (i.e., "true" gene flow unit) identified by PopCOGenT is shown alongside its pseudo-gene flow unit identified using the pseudo-gene flow unit assignment method implemented in BTyper3 v. 3.1.0 (pink branches and tip labels), with background pseudo-gene flow units and true gene flow units denoted using gray tip labels and branches. Phylogenies are rooted at the midpoint, and branch lengths are reported in substitutions per site. Genomospecies and pseudo-gene flow units were assigned using BTyper3 v. 3.1.0 and FastANI v. 1.0. Core SNPs were identified among all high-quality genomes assigned to each species using kSNP3 v. 3.92 and the optimal *k*-mer size reported by Kchooser. IQ-TREE v. 1.5.4 was used to construct each phylogeny, using the resulting core SNP alignment and the GTR+G+ASC nucleotide substitution model.
