## Supplementary Figure S32 for "No Assembly Required: Using BTyper3 to Assess the Congruency of a Proposed Taxonomic Framework for the *Bacillus cereus* group with Historical Typing Methods"

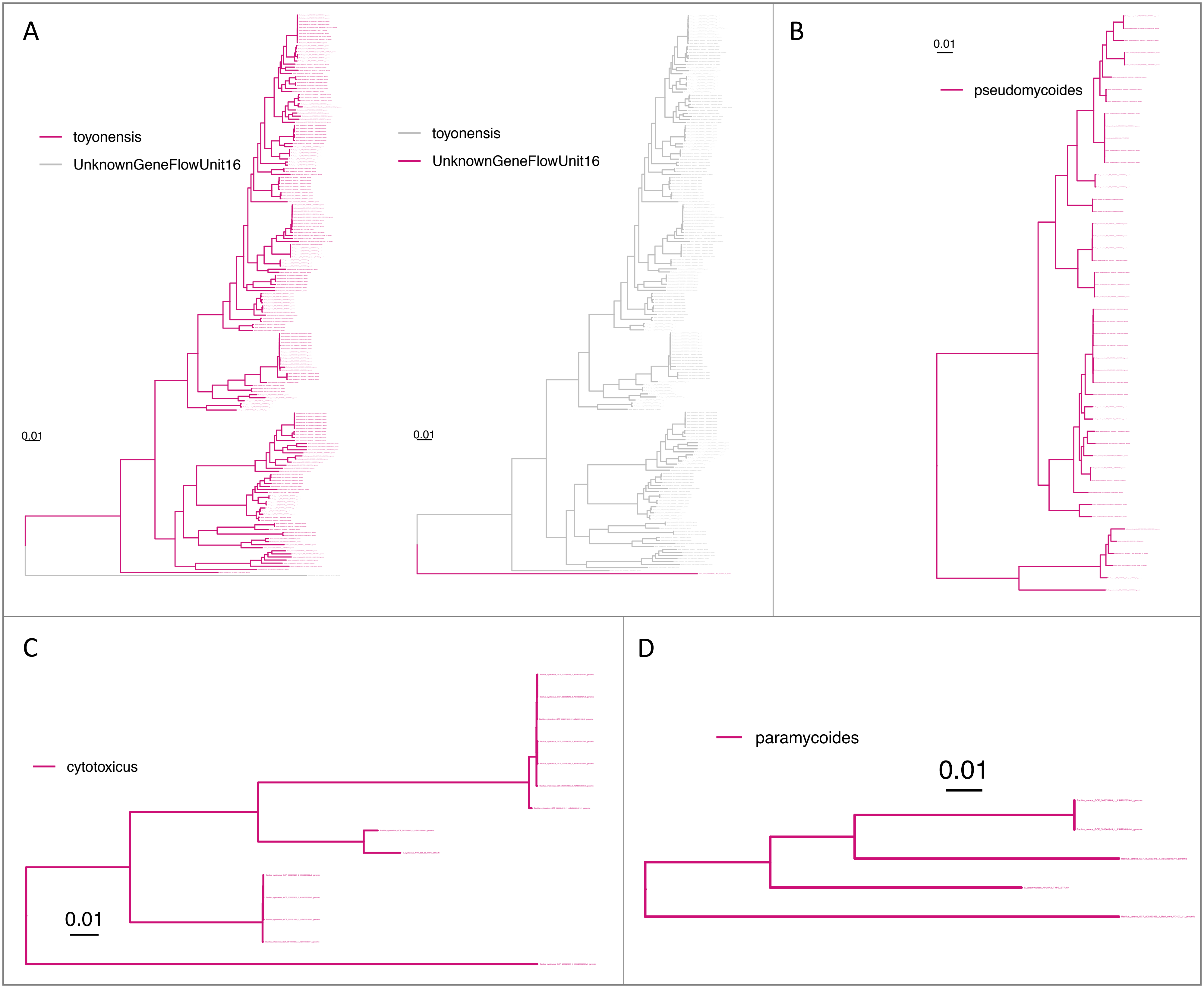


**Supplementary Figure S32.** Maximum likelihood phylogenies constructed using genome-wide core SNPs identified among all high-quality genomes assigned to the (A) *B. toyonensis*, (B) *B. pseudomycoides*, (C) *B. cytotoxicus*, and (D) *B. paramycoides* genomospecies delineated at a 92.5 average nucleotide identity (ANI) threshold. *B. luti* is not shown, as there were not enough available genomes to construct a phylogeny. Each pseudo-gene flow unit identified using the pseudo-gene flow unit assignment method implemented in BTyper3 v. 3.1.0 is shown in the foreground (pink branches and tip labels), with background pseudo-gene flow units denoted using gray tip labels and branches. Phylogenies are rooted at the midpoint, and branch lengths are reported in substitutions per site. Genomospecies and pseudo-gene flow units were assigned using BTyper3 v. 3.1.0 and FastANI v. 1.0. Core SNPs were identified among all high-quality genomes assigned to each genomospecies using kSNP3 v. 3.92 and the optimal *k*-mer size reported by Kchooser. IQ-TREE v. 1.5.4 was used to construct each phylogeny, using the resulting core SNP alignment and the GTR+G+ASC nucleotide substitution model.
